# Non-cell-autonomous promotion of pluripotency induction mediated by YAP

**DOI:** 10.1101/481127

**Authors:** Amaleah Hartman, Xiao Hu, Xinyue Chen, Anna E. Eastman, Cindy Yang, Shangqin Guo

**Affiliations:** Cell Biology Department, Yale University, New Haven, CT 06520; Yale Stem Cell Center, Yale University, New Haven, CT 06520

## Abstract

While Yes-associated protein (YAP) antagonizes pluripotency during early embryogenesis, it has also been shown to promote stemness of multiple stem cell types, including pluripotent stem cells. Whether cellular context underlies these distinct functions of YAP in pluripotency remains unclear. Here, we establish that depending on the specific cells in which it is expressed, YAP exhibits opposing effects on pluripotency induction from somatic cells. Specifically, YAP inhibits pluripotency induction cell-autonomously but promotes it non-cell-autonomously. For its non-cell-autonomous role, YAP alters the expression of many secreted and matricellular proteins including CYR61, which recapitulates the promotional effect when added as a recombinant protein. Thus, we define a unique YAP-driven non-cell-autonomous process that enhances pluripotency induction. Our work highlights the importance of considering the distinct contributions from heterologous cell types in deciphering the mechanism of cell fate control and calls for careful re-examination of the co-existing bystander cells in complex cultures or tissues.

## INTRODUCTION

Cell fate decisions are instructed by their microenvironment. A major mediator of microenvironmental signaling is the transcriptional co-activator Yes-associated protein (YAP). Diverse upstream inputs, including cell culture density and extracellular soluble factors, as well as local extracellular matrix composition, converge to regulate YAP’s nuclear entry and transcriptional activity (Azzolin et al., 2014; Dupont et al., 2011; Halder et al., 2012; Park et al., 2015; Piccolo et al., 2013; Piccolo et al., 2014; Yu et al., 2012; Zhao et al., 2010; Zhao et al., 2007). These same microenvironmental cues also profoundly influence cell fate (Chen et al., 1997; Dupont et al., 2011; Engler et al., 2006; Gilbert et al., 2010; Mammoto and Ingber, 2010; McBeath et al., 2004; Schwartz, 2010; Swift et al., 2013; Vogel and Sheetz, 2006). Accordingly, a large body of work has examined YAP’s role in cell fate decisions in various biological contexts via its direct engagement with the chromatin cell-autonomously (Barry et al., 2013; Camargo et al., 2007; Panciera et al., 2016; Qin et al., 2016; Schlegelmilch et al., 2011; Su et al., 2015; Totaro et al., 2017; Zanconato et al., 2016). However, since many of YAP’s target gene products localize outside of the cell (Katsube et al., 2009; Zhang et al., 2009; Zuo et al., 2010), it is possible that YAP influences cell fate non-cell-autonomously, a scenario that is under explored.

YAP’s role appears complex and controversial in the regulation of pluripotency. YAP functionally antagonizes pluripotency during early mouse embryonic development, when the trophectoderm fate is specified versus the pluripotent inner cell mass (ICM) (Nishioka et al., 2009; Nishioka et al., 2008). YAP is nuclear in the trophectoderm but remains cytoplasmic in the ICM, indicative of low or absent YAP transcriptional activity in the developmentally pluripotent cells of the ICM. Furthermore, YAP mRNA injected into early blastocysts direct such cells toward trophectoderm fate (Nishioka et al., 2009). These findings contrast those supporting YAP’s role in promoting pluripotency, either during pluripotency induction from fibroblasts or in pluripotency maintenance (Lian et al., 2010). In yet other studies, YAP appeared dispensable for pluripotency (Azzolin et al., 2014; Chung et al., 2016). Some of these conflicting behaviors could potentially be related to YAP’s versatile interaction with components of the β-catenin or SMAD signaling pathways (Beyer et al., 2013; Papaspyropoulos et al., 2018; Zhou et al., 2017). Overall, these studies have all focused on YAP’s cell-autonomous role in pluripotency. Heretofore, it has been unclear whether YAP functions non-cell-autonomously in pluripotency, and if so, whether such a mechanism contributes to the conflicting observations in the literature.

Pluripotency is subject to non-cell-autonomous regulation. One prominent example is the use of mitotically-inactivated “feeder” cells, which secret leukemia inhibitor factor (LIF) (Smith and Hooper, 1987; Smith and Hooper, 1983), among other potentially unidentified signals, for routine pluripotent stem cell cultivation. Additionally, the importance of non-cell-autonomous regulation is highlighted by the recent discoveries that interleukin 6 (IL-6) secreted from nearby senescent or injured cells mediates *in vivo* pluripotency induction (Chiche et al., 2017; Mosteiro et al., 2016; Mosteiro et al., 2018). Moreover, cell-plating density profoundly affects the efficiency of pluripotency induction, although the tested cell densities in the literature differ significantly (Stadtfeld et al., 2010; Wernig et al., 2008). Thus, the fate of pluripotent cells is often supported or modified by other co-existing cellular entities. Notably, cultures supporting the somatic-to-pluripotent fate conversion comprise a minority of cells that successfully acquire pluripotency and a great majority of cells that fail to do so; however, the potential contribution by the latter cells to the emerging pluripotent fate has been overlooked.

Here, we report that YAP executes two distinct functions in pluripotency induction from mouse somatic cells. While YAP potently inhibits the emergence of pluripotency cell-autonomously, it promotes pluripotency induction from nearby cells *non-cell-autonomously*. YAP-mediated non-cell-autonomous promotion does not require direct cell-cell contact, and can be recapitulated with medium conditioned by YAP-expressing cells. Further, YAP overexpression reprograms the gene expression of many secreted proteins; one of which, *Cyr61*, encoding a known matricellular protein, promotes pluripotency induction when added directly as a recombinant protein. Thus, our work elucidates a novel function of YAP that reconciles the apparent discrepancies regarding its role in pluripotency. This non-cell-autonomous function of YAP calls for careful evaluation of the role of YAP in other cellular systems.

## RESULTS

### YAP inhibits pluripotency induction cell-autonomously

Previous reports of YAP promoting pluripotency induction utilized co-transduction of viral constructs encoding YAP and the reprogramming transcription factors (Lian et al., 2010; Qin et al., 2016), yielding a mixture of cells expressing either YAP or reprogramming factors, together with cells that express both or neither, confounding the interpretation of the specific mode of YAP’s action. To dissect the cell-autonomous effect of YAP in pluripotency induction, we transduced either wildtype (wtYAP) or constitutively-active YAP (caYAP, in which two inhibitory phosphorylation sites are mutated (Zhao et al., 2010; Zhao et al., 2007)), alongside a control empty vector (EV), into transgenic reprogrammable MEFs which express a polycistronic cassette encoding Oct4, Klf4, Sox2 and cMyc (OKSM) upon Doxycycline (Dox) treatment (Stadtfeld et al., 2010) (Fig 1A-B). Successfully transduced cells were FACS-sorted based on their expression of mCherry, also encoded by the vectors (Fig. 1A), and replated onto mitotically-inactivated feeder MEFs for further reprogramming. This approach ensures that most or all cells express both YAP and OKSM. Mature iPSCs were identified by their expression of GFP from the endogenous Oct4 locus (Stadtfeld et al., 2010) (Fig 1C-D). Reprogrammable MEFs co-expressing wtYAP or caYAP produced significantly fewer Oct4:GFP+ iPSCs compared to EV control (Fig 1E-F). This reduction could not be accounted for by increased vector silencing or compromised viability/proliferation by YAP-expressing cells, as mCherry+ cells were abundant in YAP-expressing cultures, although such cells were negative for Oct4:GFP (Fig 1C-D). Even after prolonged exposure to both OKSM and YAP expression (75 days), the cells remained negative for Oct4:GFP (Fig 1G). Furthermore, rather than compact dome-shaped colonies characteristic of mouse pluripotency, wtYAP-expressing cells produced colonies with flat morphology (Fig 1G), while caYAP-expressing cells did not withstand further passaging beyond approximately 28 days (data not shown). The wtYAP-overexpressing flat colonies could be passaged indefinitely in mouse embryonic stem cells (mESC) maintenance conditions, although they were not pluripotent as determined by the lack of endogenous pluripotency gene expression (Fig 1H-I and Fig S1A-C). Intriguingly, following long term culture in Dox, a small subset of the YAP-transduced cells emerged as mCherry+Oct4:GFP+ (Fig S1A,B). However, these cells no longer displayed elevated YAP or target gene expression (Fig S1D), suggesting that either mature pluripotent cells could dampen YAP activity or that low YAP-activity cells were advantageous during prolonged culture. In conclusion, MEFs simultaneously over-expressing YAP and OKSM failed to establish pluripotency.

**Figure 1:**
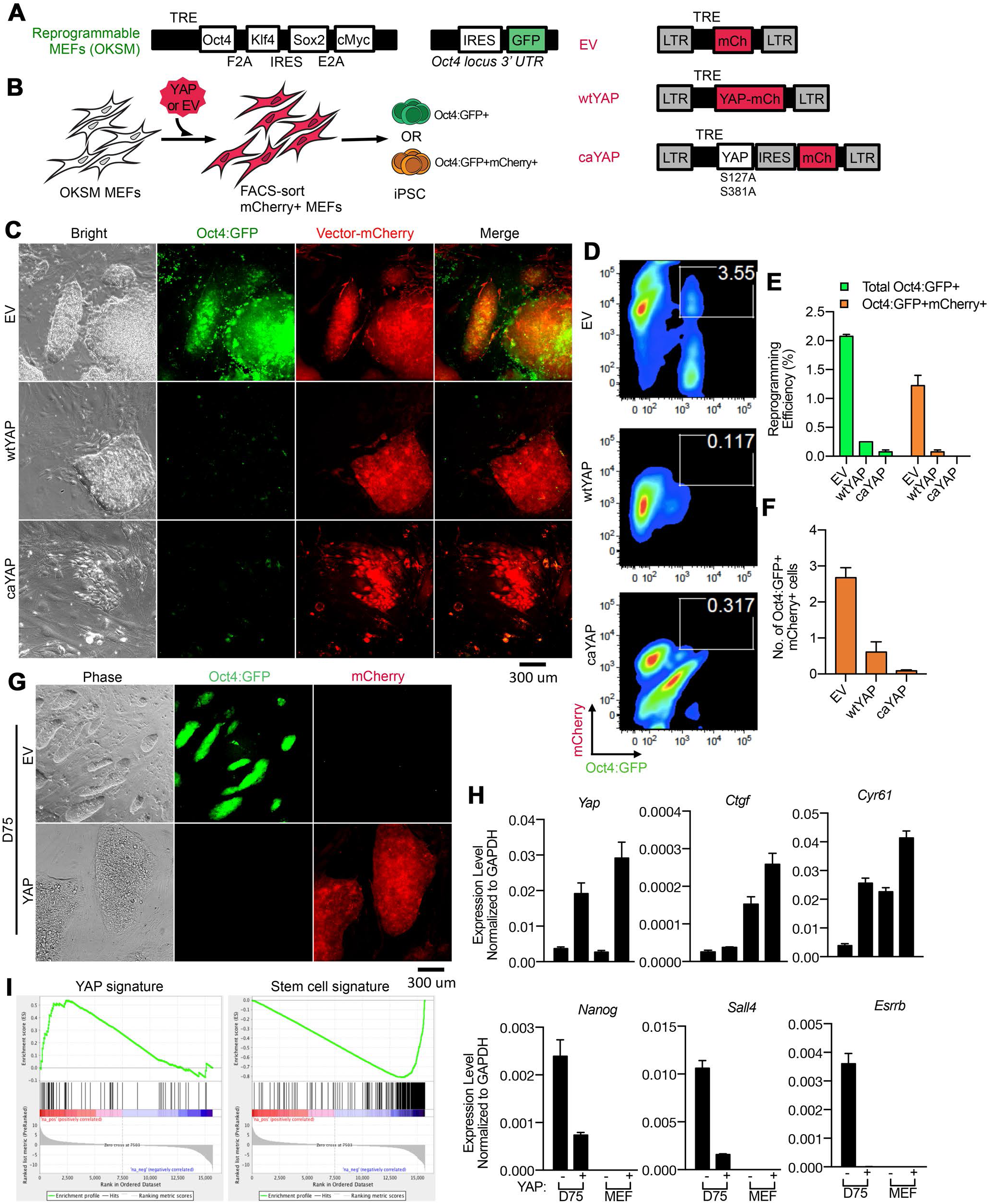
YAP inhibits pluripotency induction cell autonomously. (A) Top: Transgenic reprogrammable system for OKSM expression under the control of a tetracycline-responsive element (TRE). These cells also express a GFP reporter under the control of the endogenous Oct4 promoter. Right: Lentiviral vectors encoding mCherry (empty vector; EV), wildtype YAP fused to mCherry (wtYAP), or constitutively active YAP followed by an internal ribosome entry site (IRES) and mCherry (caYAP). LTR: long terminal repeats. (B) Experimental scheme illustrating primary OKSM-expressing MEFs transduced with viral vectors in (A), FAGS-sorted on day 3 of Dox treatment based on mCherrypositivity and replated to allow further reprogramming. The expression status of Oct4:GFP was determined in the resulting iPSCs in relation to their expression of mCherry, shown in (C-D). (C) Representative images of reprogramming cultures after 15 days of Dox treatment. (D) Representative FACS plots of reprogramming cultures after 20 days of Dox treatment. Gated population denotes Oct4:GFP and mCherry double-positive cells. (E) Quantification of reprogramming efficiency based on the number of Oct4:GFP+ colonies and the number of Oct4:GFP and mCherry double-positive colonies. (F) Quantification of the absolute numbers of Oct4:GFP and mCherry double-positive cells in each culture condition (total cell number multiplied by % double-positive cells in (D)). (G) Representative colony morphology after long-term culture (Day 75) in mESC conditions derived from OKSM MEFs expressing control (EV) or wildtype YAP. (H) RT-qPCR analysis of gene expression in cells shown in (G), with MEFs expressing mCherry alone (EV, denoted as YAP−) or wtYAP (YAP+) as controls. (I) Differentially expressed genes from mRNA-seq analysis between cells shown in (G) by gene set enrichment analysis. YAP signature is from (Zhang et al., 2009) and stem cell signature is from Cluster Ill (Polo et al., 2012).

To examine YAP’s cell-autonomous effect on pluripotency induction from other somatic cell types, we expressed YAP in reprogrammable granulocyte-monocyte progenitors (GMPs) (Fig S2). Similar to MEFs, YAP co-expression inhibited GMP reprogramming (Fig S2B-C). Of the Oct4:GFP+ cells that arose from YAP-transduced cultures, the percentage of Oct4:GFP+ cells decreased upon further culture, while the EV-transduced cultures behaved in the opposite manner (Fig S2D-E), suggesting a competitive disadvantage for Oct4:GFP+ cells initially co-expressing YAP. Furthermore, the fluorescence intensity of Oct4:GFP was lower in YAP co-expressing cells (Fig S2F), suggesting partial activation of the endogenous Oct4 locus. Taken together, these results further support that YAP compromises pluripotency induction when co-expressed with the reprogramming factors.

The inhibition of pluripotency induction by co-expressed YAP prompted us to examine the behavior of endogenous YAP during reprogramming. We assessed the subcellular localization of endogenous YAP in MEFs transduced with polycistronic expression of OSKM and mCherry (Fig 2A). By comparing OSKM-expressing (mCherry+) and wildtype (mCherry−) cells of the same culture, the effect of cell density or other culture-related variables are minimized, allowing for examination of YAP localization largely consequent to reprogramming factor expression. On day 4, the OSKM-expressing (mCherry+) cells displayed significantly lower nuclear YAP signal than the wildtype (mCherry−) cells of the same culture (Fig 2B-C). These results suggest that cells of lower YAP activity are more permissive of OSKM expression, or that OSKM expression actively antagonizes YAP nuclear localization. Consistently, the expression of YAP target genes (*Ctgf* and *Cyr61*) was diminished in mCherry+ cells as reprogramming proceeded (Fig 2D). Reduced YAP target gene expression was also evident in cells previously shown to have enhanced reprogramming capacity (Fig 2E) (Guo et al., 2014). Finally, the expression of YAP target genes was lower in iPSCs and ESCs as compared to MEFs, suggesting reduced YAP activity in pluripotent stem cells (Fig 2F). This could be the result of active downregulation of YAP activity in pluripotency, or a consequence of the drastic morphological changes accompanying the acquisition of pluripotency. Nevertheless, pluripotency coincided with low levels of endogenous YAP target gene expression, supporting YAP’s inhibitory role in pluripotency induction cell-autonomously.

**Figure 2:**
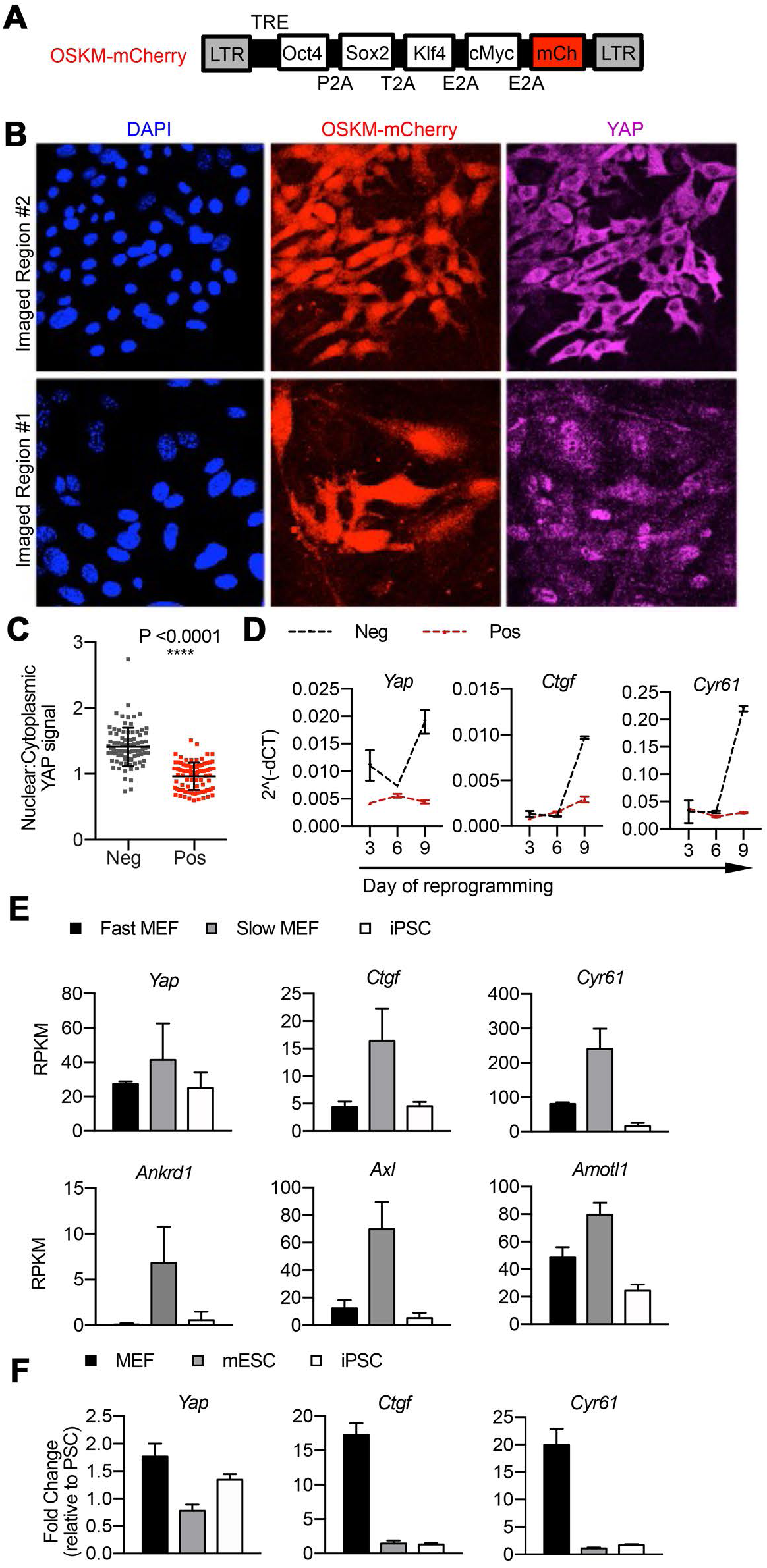
Reduced endogenous YAP target gene expression correlates with the induction of pluripotency. (A) Schema of polycistronic lentiviral vector that encodes OSKM-mCherry. (B) Primary MEFs were transduced with viral OSKM-mcherry shown in (A) and treated with Dox for four days, and fixed and st in d with a YAP-specifiantibody. Two representative regions within the same culture of distinct local cell density are shown. (C) Quantification of the endogenuous nuclear and cytoplasmic YAP signal comparing the mCherry- (Neg) and mCherry+ (Pos) cells. Nuclear are was defined by DAPI+ region; cytoplasm area was defined by a pan-cellular GFP (not shown) minus the DAPI+ region. Each dot donates the nuclear to cytoplasmic YAP sugnal of a single cell (N = 100 each). (D) RT-qpcr analyses for endogenous Yap and YAP target genes Ctgf and Cyr61 in mCherry- (Neg) and mCherry+ (Pos) cells sorted after 3, 6 or 9 days of Dox treatment. (E) Expression of endogenous Yap and YAP target genes *Ctgf*, *Cyr61*, *Ankrd1*, *Axl* and *Amot1* by mRNA-seq in fast-cycling cells that exhibit enhanced reprogramming efficiency compared to slow-cycling cells and mature iPSCs. Data are re-plotted from (Guo et al., 2014). RPKM: reads per kilobase of transcript per million mapped reads. (F) RT-qPCR of endogenous *Yap* and YAP target genes *Ctgf* and *Cyr61* in primary MEFs, mESCs and iPSCs.

### Ectopic YAP expression does not promote pluripotency maintenance

Because YAP has been previously reported to promote pluripotency maintenance (Lian et al., 2010; Qin et al., 2016), we considered the possibility that YAP promotes the maintenance of established pluripotency, even though it did not promote the somatic-to-pluripotency transition. To examine a cell-autonomous effect of YAP on established pluripotency, we transduced wtYAP into mESCs harboring an Oct4:GFP reporter (Fig 3A). Transduced cells (mCherry+) were FACS-sorted and replated in mESC maintenance conditions (Fig 3B). Although the total number of Oct4:GFP+ colonies was similar between EV and YAP transduced cultures, the YAP-transduced ESC cultures had significantly fewer colonies that were mCherry+ (Fig 3C). This could suggest a competitive disadvantage of the YAP-expressing mESCs, or that the YAP-expressing mESCs silenced the viral vector more effectively. In either case, YAP expression did not favor the maintenance of established pluripotency, at least under standard mESC culture conditions.

**Figure 3:**
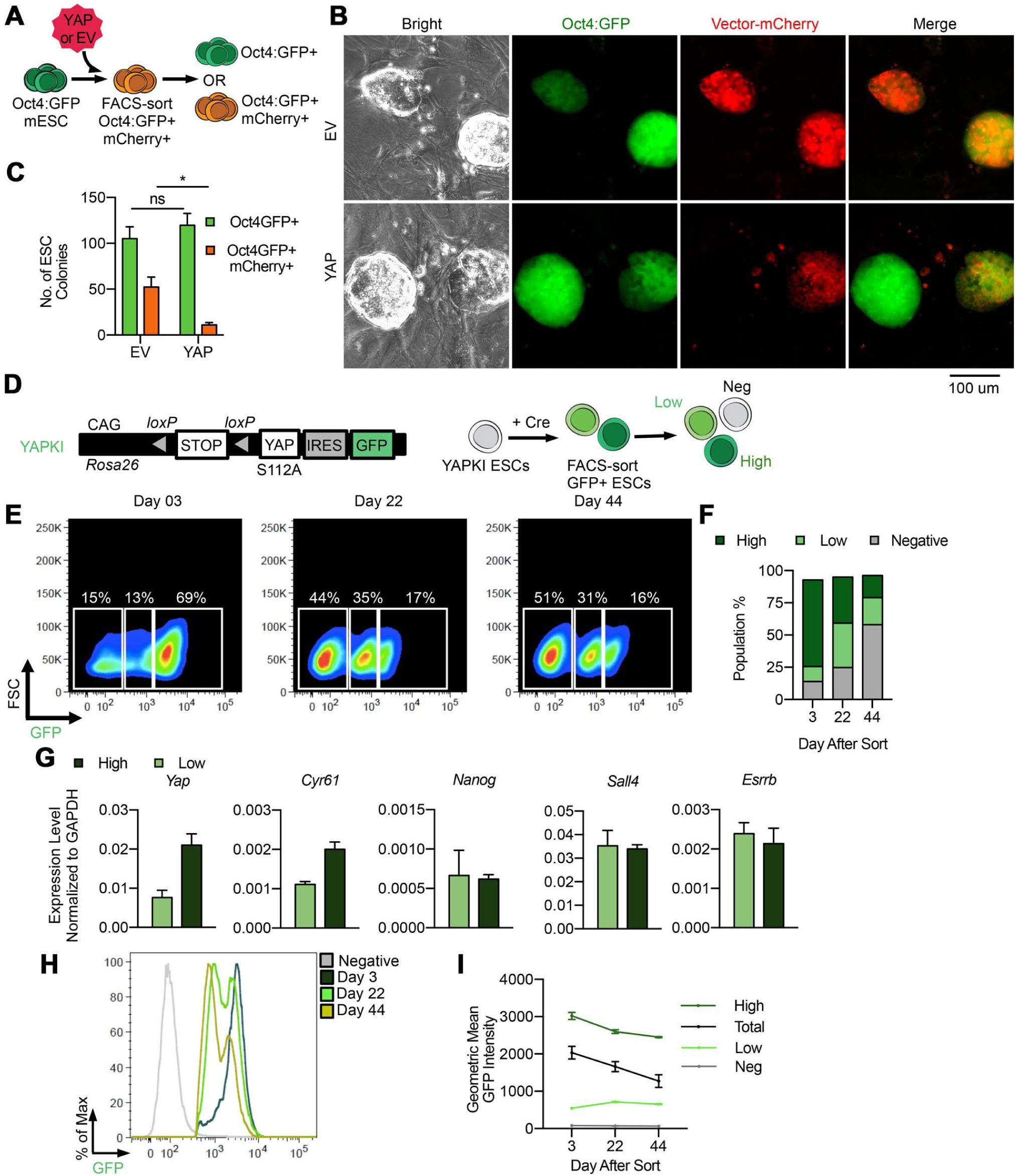
Ectopic YAP expression does not promote pluripotency maintenance. (A) Experimental scheme illustrating mESCs expressing an Oct4:GFP reporter transduced with EV or wtYAP (same as in Fig 1A) and, FACS-sorted for Oct4:GFP and mCherry double-positive cells. The expression status of mCherry was determined in the resulting mESCs, shown in (B-C). (B) Representative images of Oct4:GFP mESCs and their mCherry expression after 7 days of culture. (C) Quantification of the total number of Oct4:GFP+ mESC colonies and those that co-expressed mCherry. (D) Left: Schema of YAPKI allele in the *Rosa26* locus, containing a coding sequence for caYAP and IRES-GFP preceded by a *loxP*-flanked STOP signal. Right: mESCs harboring this allele undergo Cre-mediated recombination, yielding two populations of GFP+ cells, which were FACS-sorted and cultured in standard mESC maintenance condition. The relative abundance of the GFP-negative, GFP-low, and GFP-high populations was analyzed over time, shown in (E-I). (E) Representative FACS plots of the GFP-negative, GFP-low and GFP-high populations on day 3, 22 and 44 after sorting. (F) Quantification of data shown in (E). (G) RT-qPCR analyses of endogenous *Yap*, *Cyr61*, and pluripotency network genes *Nanog*, *Sall4* and *Esrrb* in the GFP-high (High) and GFP-low (Low) mESCs after 3 days in culture following FACS-sort. (H) Representative FACS plot showing the evolution of fluorescence intensity of the GFP+ population over time. (I) Quantification of GFP geometric mean fluorescence intensity in the total GFP+ population (Total), GFP-low (Low), GFP-high (High) and GFP-negative (Negative) populations over time.

To circumvent silencing of the virally expressed YAP, we induced ectopic YAP expression in mESCs containing a loxP-STOP-loxP-caYAP(S112A)-IRES-GFP cassette (YAPKI mESCs) in the *Rosa26* locus (Su et al., 2015). We transduced the YAPKI mESCs with a lentiviral Cre, whose transient activity permanently activated the YAPKI allele as well as the GFP reporter (Fig 3D). Shortly after Cre transduction, cells that successfully underwent recombination were FACS-sorted and replated in maintenance conditions (Fig 3D). Consistent with the original report describing the YAPKI allele (Su et al., 2015), two populations of GFP+ cells with distinct intensities emerged after recombination (Fig 3E-F), with only the GFP-high cells expressing increased YAP target genes (Su et al., 2015). We confirmed that the GFP-high mESCs indeed exhibited elevated *Yap* and target gene expression, while expressing comparable levels of pluripotency genes to the control GFP-low cells (Fig 3G). We observed that the percentage of GFP-high cells decreased relative to the GFP-low cells in the same culture over time (Fig 3E-F and Fig 3H, S3A-B). A substantial portion of the culture became negative for GFP, likely due to the expansion of the few GFP-cells from the original sorting (Fig 3E-F). Further, even among cells that remained within the GFP-high gate, their GFP intensity decreased over time (Fig. 3I, S3C). Taken together, these data indicate that ectopic YAP expression does not promote, but rather is competitively unfavorable for, pluripotency maintenance when expressed cell-autonomously.

### YAP promotes pluripotency induction in a non-cell-autonomous manner

Having ruled out a cell-autonomous effect by YAP in pluripotency induction or maintenance, we examined whether YAP regulates pluripotency non-cell-autonomously. We delivered the reprogramming factors via lentivirus into MEFs to crudely establish a heterogeneous cell culture comprised of cells expressing either OSKM, YAP, both, or neither (Fig 4A), similar to the previous studies (Lian et al., 2010; Qin et al., 2016). We tagged YAP and OSKM-expressing vectors with fluorescent markers – GFP and mCherry, respectively (Fig 4A), to track the contribution of heterogeneous cell types co-existing in the co-transduced cultures (Fig 4B-C). Consistent with the previous report (Lian et al., 2010), we observed a two-fold increase in the total number of colonies when wtYAP was co-transduced (Fig 4D). While more colonies were present in the wtYAP co-transduced cultures (Fig 4D), few of them co-expressed YAP as indicated by their lack of GFP (i.e. YAP) expression (Fig 4C-E). Interestingly, caYAP co-transduction reduced the number of total colonies (Fig 4E). The absence of YAP+ colonies could not be simply accounted for by enhanced vector silencing, as GFP+ cells were still abundant at this time point even though they did not appear as colonies (Fig 4C). Assessing the activation of endogenous pluripotency by immunofluorescence staining of Nanog yielded similar results (Fig 4F-H). Overall, these data demonstrate that while co-transduced wtYAP promotes the emergence of iPSC colonies, the colonies themselves do not co-express YAP. The presence of non-pluripotent YAP-expressing fibroblasts strongly suggests a non-cell-autonomous effect mediated by YAP. The fact that promotion was only observed with co-transduced wtYAP, but not caYAP, suggests that the non-cell-autonomous effect is limited to wtYAP, while the cell-autonomous inhibition is shared by both.

**Figure 4:**
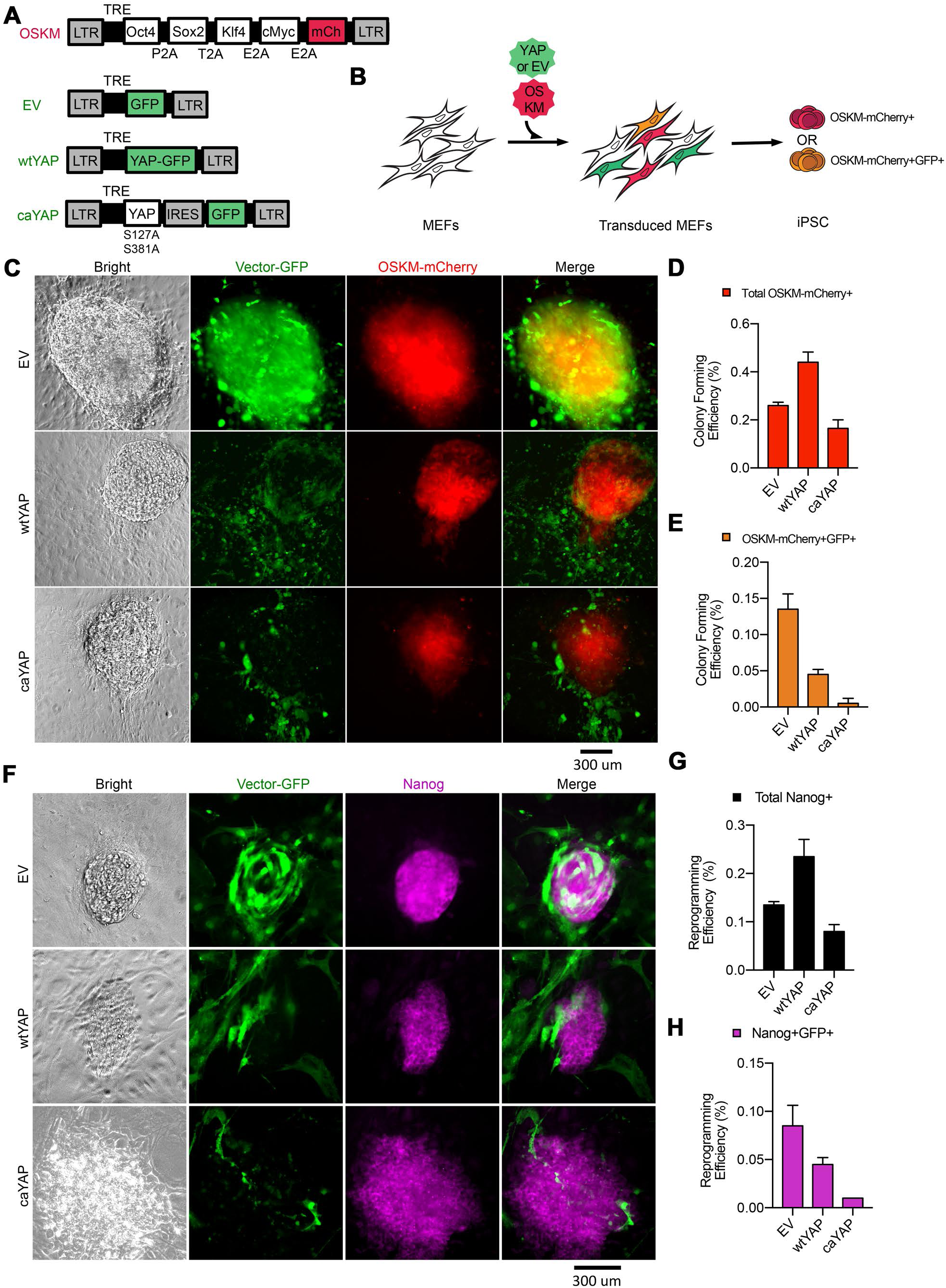
YAP promotes pluripotency induction in mixed cultures. (A) Schema of lentiviral vectors used to generate a mixture of cells that express OSKM, YAP, both or neither. (B) Experimental scheme illustrating the heterogeneous cell types co-existing in culture. The expression status of GFP from EV or YAP-expressing vectors as determined in the resulting colonies, shown in (C-H). (C) Representative OSKM-mCherry+ colony images after 18 days of reprogramming from MEF cultures co-transduced with EV, wtYAP, or caYAP, each of which also expresses a GFP reporter. (D) Quantification of total OSKM-mCherry+ colonies in (C). Representative microscope images of resulting colonies after 18 days of reprogramming derived from each co-transduced population. (E) Quantification of the OSKM-mCherry+ colonies also positive for GFP in (C). (F) Representative images of iPSC colonies after 18 days of reprogramming immuno-stained for the endogenous Nanog protein. (G) Quantification of total Nanog+ iPSC colonies in (F). (H) Quantification of Nanog+ iPSC colonies also positive for GFP in (F).

To directly test the possibility of a non-cell-autonomous role, we mixed, in a controlled manner, two types of cells: one expressing wtYAP-GFP (YAP) or a control GFP empty vector (EV) and the other expressing OSKM-mCherry, with GFP+ cells serving as the “feeder” cell population (Fig 5A). After FACS-sorting of each population, cells were replated together at varying ratios of feeder-to-reprogramming cells, while maintaining overall cell-plating density. Strikingly, all conditions in which reprogramming cells were co-cultured with YAP-feeders produced more colonies (Fig 5B-C). The promotional effect became more pronounced as the ratio of YAP-feeders increased (Fig 5B-C). These data demonstrate that YAP-expressing fibroblasts promote pluripotency induction non-cell-autonomously.

**Figure 5:**
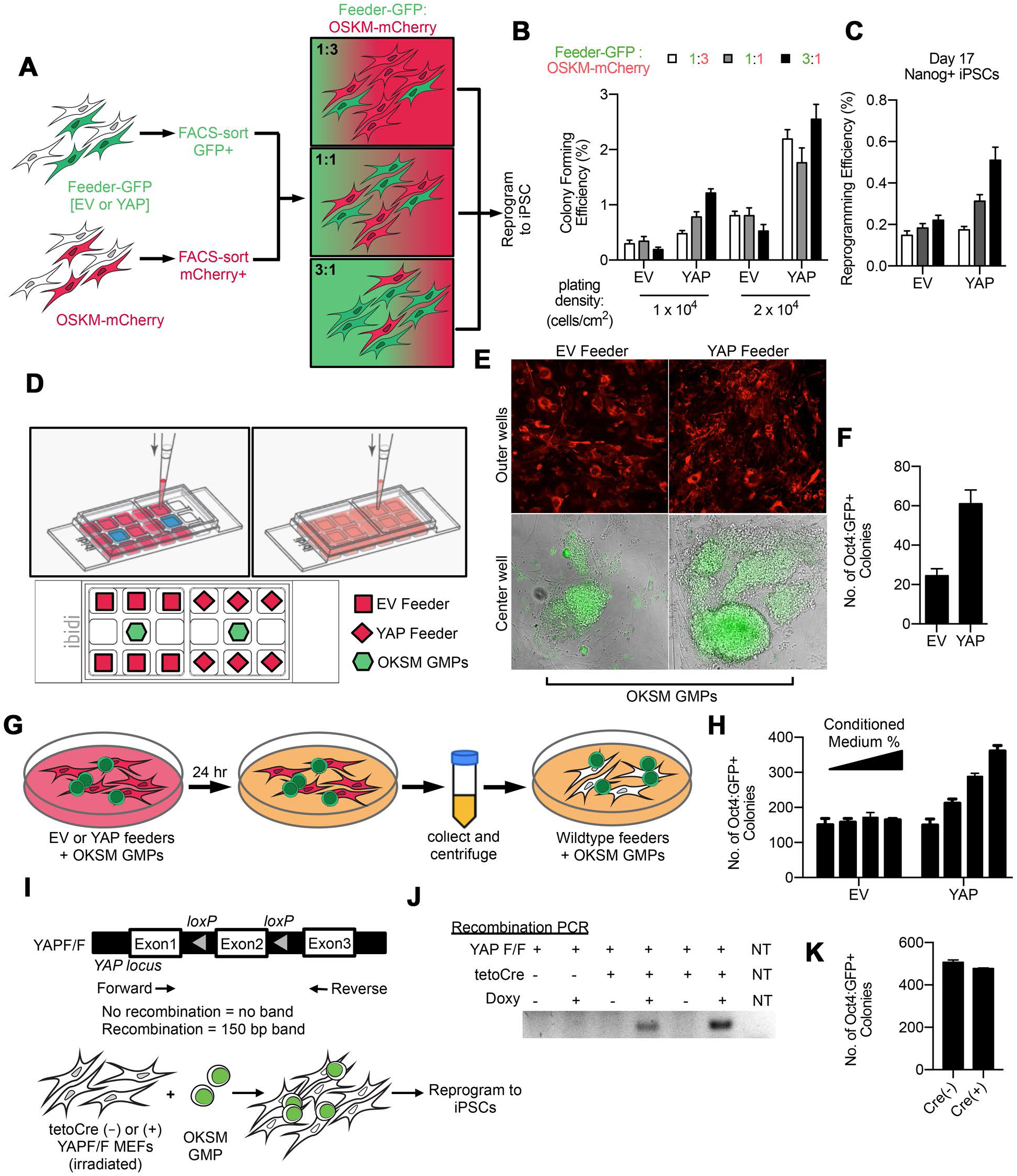
YAP promotes pluripotency induction cell non-autonomously. (A) Experimental scheme illustrating the mixing of two separate populations of primary MEFs: one expressing GFP (EV) or wildtype YAP fused to GFP (YAP) corresponding to the “feeder” cells and the other expressing OSKM-mCherry. The cells were FACS sorted and replated at varying ratios as indicated, at two plating densities (1 × 10^4^ and 2 × 10^4^ cells/cm^2^). Resulting iPSC colonies were scored on day 15-17 as shown in (B-C). (B) Quantification of colony forming efficiency (mCherry+) on day 15 of reprogramming. (C) Quantification of Nanog+ iPSC colonies on day 17, from cultures plated at 1 × 10^4^ cells/cm^2^. (D) Top: Schema illustrating a co-culture device that allows physically separated cells to share medium. Each of two major wells has 9 minor wells. Upon seeding cells in the individual minor wells, the major well was filled with medium so that the medium was shared among cells across the 9-minor-well unit. Bottom: Schema illustrating the co-culture of mitotically-inactivated feeders expressing either control (EV) or wildtype YAP fused to mCherry (YAP) in outer minor wells and reprogrammable GMPs in the central minor well. (E) Representative images of Oct4:GFP+ iPSCs (bottom) formed in the center well, fed by either EV-feeders or YAP-feeders, on day 5 of reprogramming. (F) Quantification of data shown in (E). (G) Experimental scheme illustrating the preparation of conditioned medium. (H) Quantification of Oct4:GFP+ iPSC colonies cultured with varying proportion of conditioned medium on day 5 of reprogramming. (I) Top: Schema of the YAP conditional allele. Arrows denote PCR primers for detecting the recombined product. Bottom: Schema for using tetOCre- or tetOCre+ YAPF/F MEFs in supporting reprogramming of OKSM GMPs. (J) Representative PCR reactions confirming successful recombination, i.e. YAP inactivation, in Cre+ MEFs. NT: no template control. (K) Quantification of the number of Oct4:GFP+ iPSC colonies by OKSM GMPs on tetOCre- or tetOCre+ YAPF/F MEFs on day 5 of reprogramming.

To determine if the promoting effect requires direct cell-cell contact, we utilized a co-culture device in which different cell types are kept physically separate, while sharing the same medium (Fig 5D). Reprogrammable GMPs were plated in the center minor well within each co-culture unit to be “;fed”; by mitotically-inactivated feeder MEFs expressing either EV (mCherry) or YAP (YAP-mCherry) in the surrounding outer minor wells (Fig 5D). For experiments using this co-culture device, we decided to test GMPs in the center well because these wells had limited growth area and were not amenable to accommodating the extensive cell proliferation required for MEF reprogramming. The OKSM GMPs sharing medium with YAP-feeders yielded more Oct4:GFP+ colonies compared to those sharing medium with EV-feeders (Fig 5E-F). Thus, direct cell-cell contact is not necessary for YAP to promote pluripotency induction. To directly test whether conditioned medium is sufficient to mediate the YAP-feeder effect, we compared the reprogramming efficiency of reprogrammable GMPs fed by medium conditioned by YAP-feeders or EV-feeders (Fig 5G-H). More Oct4:GFP+ iPSC colonies arose when cultured in YAP-feeder conditioned medium. These data demonstrate that YAP promotes pluripotency induction non-cell-autonomously, which is at least partly mediated by components that exist in the medium.

To assess whether YAP is required for the feeder cells to support pluripotency induction, we derived MEFs carrying a conditional YAP allele (YAP F/F) (Su et al., 2015) crossed with a Dox-inducible Cre (tetOCre) (Perl et al., 2002); MEFs derived from the same litter but negative for Cre were used as control (Fig 5I). Following 2 days of Dox treatment to ensure recombination of the YAP F/F allele (Fig 5J), reprogrammable GMPs were plated onto the feeder cells that were either positive or negative for tetOCre, in the continued presence of Dox. YAP deletion in the feeder cells did not adversely affect the number, size or morphology of the emerging iPSCs (Fig 5K, and data not shown). Thus, while YAP gain-of-function in the feeder cells promoted pluripotency induction, YAP does not appear to be essential, at least in the commonly-used pluripotency culture conditions in which key growth factors such as LIF are abundant.

### YAP target CYR61 promotes pluripotency induction

For the conditioned medium to mediate more effective reprogramming, it could either contain increased levels of factor(s) that promote pluripotency induction or reduced levels of factor(s) that inhibit the process. To uncover the identity of potential secreted factor(s), we first performed cytokine/growth factor detection arrays using medium conditioned by YAP-feeders or EV-feeders (Fig S4A-B). Only three out of 111 probed proteins were differentially present (at least two-fold difference) in two independent experiments: Pentraxin-3 (PTX-3), CCL6/C10, and CCL11 (Fig S4A-B), with PTX-3 being the only protein increased in the YAP-feeder conditioned medium (Fig S4A-B).

To look for secreted protein factors beyond those on the growth factor array, we carried out mRNA-seq comparing MEFs transduced with EV or wtYAP and FACS-sorted based on their encoded fluorescence reporter (mCherry or GFP) (Fig S4C). Among the up-regulated genes, known YAP-target genes *Cyr61* and *Ctgf* were both increased in the YAP-overexpressing samples (Fig S4D), supporting effective YAP-overexpression and transcriptional activity. The overall gene expression was similar between EV- and YAP-expressing cells, i.e., the number of differentially expressed genes was low (Fig S4E-F). Interestingly, among the differentially expressed genes, many belonged to the “;extracellular region”;, “;extracellular exosome”; or “;extracellular matrix”; (Fig S4G-H). This was the case for both the up-regulated and down-regulated genes. These data indicate that YAP expression potentially alters the compositions of the microenvironment, providing a molecular basis for a non-cell-autonomous role for YAP.

Given the recent discovery of IL-6 in promoting pluripotency induction (Brady et al., 2013; Mosteiro et al., 2018), we focused on the up-regulated genes in YAP-expressing cells. Analysis of the differentially expressed genes revealed 40 that encode secreted proteins with increased expression in YAP-expressing MEFs (Fig S4K). Specifically, we chose to test whether recombinant CTGF, CYR61 or PTX3 promote reprogramming, using IL-6 as a positive control (Brady et al., 2013; Mosteiro et al., 2018). As expected, recombinant IL-6 resulted in ∼2-fold increase in the number of Oct4:GFP+ cells and colonies from reprogrammable MEFs (Fig 6A-D) or GMPs (Fig S5A). While recombinant CTGF and PTX3 had no effect on reprogramming (Fig 6A-D, Fig S5B-D), recombinant CYR61 promoted reprogramming from MEFs, assessed by the percentage of Oct4:GFP+ cells (Fig 6A,C) or by the number of Oct4:GFP+ iPSC colonies (Fig 6B,D). The extent of promotion was similar to that of recombinant IL6 (Brady et al., 2013). Furthermore, recombinant CYR61 also promoted reprogramming starting from GMPs (Fig S5A).

**Figure 6:**
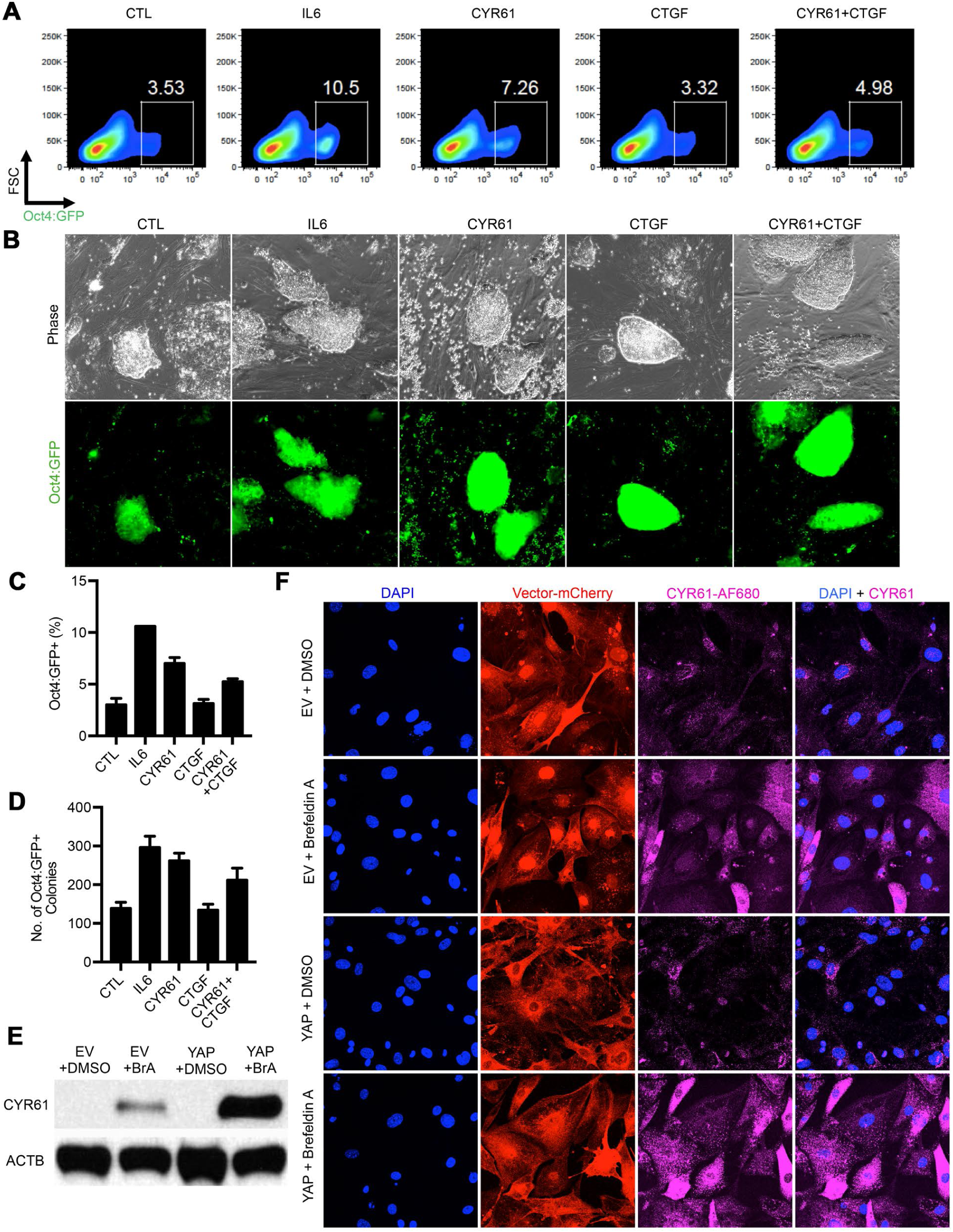
Recombinant CYR61 protein promotes pluripotency induction. (A) Representative FAGS plots for Oct4:GFP+ cells from reprogrammable MEFs in the presence of recombinant proteins IL6, CYR61, CTGF, or both CYR61 and CTGF on day 15 of reprogramming. (B) Representative images of the resulting iPSCs colonies, following the same treatment shown for (A). (C) Quantification of the percent of Oct4:GFP+ cells from (A). (D) Quantification of the number of Oct4:GFP+ iPSCs in (B). (E) lmmunoblot for CYR61 in EV and YAP feeders, treated overnight with DMSO or 0.5 ng/ml Brefeldin-A in DMSO (BrA), with fl-actin (ACTB) as a loading control. (F) lmmunofluorescence for CYR61 in EV and YAP feeders, treated overnight with DMSO or 0.5 ng/ml Brefeldin-A in DMSO (BrA).

Lastly, we validated the increased production and secretion of CYR61 protein (Fig 6E-F) by immunofluorescence and immunoblot. We treated EV- and YAP-expressing feeder MEFs with Brefeldin-A, which inhibits the protein secretory pathway (Sciaky et al., 1997). Compared to vehicle DMSO-treated cells, Brefeldin-A treatment increased CYR61 signals in both EV- and YAP-expressing feeders (Fig 6E-F), indicating that the CYR61 protein is indeed being secreted. The intracellular accumulation of CYR61 became exaggerated in the YAP-expressing feeders (Fig 6E-F). Thus, the YAP-expressing cells can supply higher levels of CYR61 to the shared culture medium, which is sufficient to promote pluripotency induction non-autonomously.

### Cell density modulates both cell-autonomous and non-cell-autonomous functions of YAP

Since YAP’s subcellular localization and activity are regulated by cell-plating density (Dupont et al., 2011), we next tested how reprogramming efficiency responds to varying cell densities as a means to indirectly regulate YAP. We first tested how cell density modulates reprogramming (Fig S6). MEFs were transduced with viral OSKM-mCherry (Fig S6A) and the same transduced culture was trypsinized and replated at varying densities. This ensured the same percentage of cells expressing OSKM in each condition and that the only changing variable was cell-plating density (Fig S6B). Reprogramming efficiency increased with cell density as assessed by either alkaline phosphatase staining (Fig S6C) or Oct4:GFP reporter activity (Fig S6D). While the starting transduction efficiency was identical in all density conditions, the percentage of OSKM-mCherry+ cells increased with cell-plating density (Fig S6E), indicating a positive selection for OSKM-expressing cells at higher cell-plating density. Thus, higher cell plating density promotes pluripotency induction.

To test if inactivated YAP underlies the beneficial effect at high cell density, we transduced reprogrammable MEFs with YAP or control EV, both encoding mCherry reporters (Fig S6F). Successfully transduced cells were FACS-sorted and replated at varying densities to reprogram (Fig S6G). YAP overexpression cell-autonomously negated the promotional effect of high cell density, as assessed by the number of resulting Oct4:GFP+ iPSC colonies or the percentage of Oct4:GFP+ cells (Fig S6H-I). Interestingly, the fluorescence intensity of YAP-mCherry decreased at high cell density, while the mCherry intensity remained constant for the EV control (Fig S6J), potentially indicating a negative regulation of the YAP-mCherry fusion protein in dense cultures. Taken together, these data are consistent with the notion that high plating density inactivates YAP and thereby alleviates the OSKM-expressing cells of YAP’s cell-autonomous inhibition on pluripotency induction.

We also examined how cell density modulates YAP’s non-cell-autonomous effect on reprogramming. Similar to the experiments above, we mixed YAP-expressing MEFs and OSKM-expressing MEFs at varying ratios while controlling for total plating density (Fig 5A). At higher cell density, increasing YAP feeder ratio did not further boost reprogramming efficiency (Fig 5B), suggesting that the ectopically expressed wtYAP was likely inactivated by cell crowding, or alternatively, the relevant factors produced by YAP feeder cells had reached saturation.

Taken together, cell density affects both modes of YAP function. At high cell density, the non-cell-autonomous promoting effect plateaus while the cell-autonomous inhibition of pluripotency induction is diminished. Thus, controlling cell-plating density could be a simple and effective way to boost efficiency of reprogramming.

## DISCUSSION

We report that YAP has dual functions in pluripotency induction: it inhibits pluripotency induction cell-autonomously, consistent with studies of early embryogenesis (Nishioka et al., 2009; Nishioka et al., 2008), and promotes it in a non-cell-autonomous manner. Further, we demonstrate that YAP induces expression changes in many genes encoding extracellular proteins. We propose that YAP promotes pluripotency induction non-cell-autonomously by reprogramming the microenvironment, and we have identified one of YAP’s targets, the matricellular protein CYR61, in mediating the promotional effect. It remains possible that additional mechanisms mediate YAP’s non-cell-autonomous effect, such as by inducing the production of certain miRNAs, which could be secreted via extracellular vesicles such as exosomes (Chen et al., 2008; Kosaka et al., 2010; Vickers et al., 2011; Wang et al., 2010). The non-cell-autonomous mode of YAP’s action likely contributes to YAP’s seemingly conflicting roles in pluripotency. For example, in previous studies where YAP was seen to promote pluripotency induction and maintenance of mouse cells (Lian et al., 2010) or naïve pluripotency of hESCs (Qin et al., 2016), exogenous YAP was virally introduced into bulk cultures, likely creating cultures of heterogeneous YAP expression, thereby preserving the possibility that the observed YAP effects were non-cell-autonomous.

Since their isolation in 1981, pluripotent mESCs have been traditionally cultured on mitotically-inactivated “feeder” MEFs. The feeder MEFs inhibit spontaneous differentiation in a non-cell-autonomous manner. Even though feeders have been conventionally used for decades, their exact contribution to pluripotent stem cell and reprogramming cultures is often overlooked. It is known that feeder MEFs secrete LIF (Smith et al., 1988; Smith and Hooper, 1987; Smith et al., 1992; Smith and Hooper, 1983); however, the understanding of the mechanism of action of feeder MEFs in the *in vitro* PSC niche largely stops here. Our work demonstrates that within a heterogeneous reprogramming culture, the non-reprogramming cells, analogous to the LIF-secreting feeder cells, are not merely passive bystanders. Instead, they could actively participate in nearby cell fate conversion by reprogramming the microenvironment they share.

YAP transcriptionally controls *Cyr61*, which encodes a secreted matricellular protein. Functionally, CYR61 modulates inflammation and senescence (Jun and Lau, 2010), both of which have been implicated in pluripotency induction via non-cell-autonomous mechanisms. Specifically, activation of innate immunity was shown to increase reprogramming in the process of “transflammation” (Lee et al., 2012). A recent study reveals that reprogramming induced in live animals triggers senescence in some cells and reprogramming in others (Mosteiro et al., 2016; Mosteiro et al., 2018). Interestingly, within this *in vivo* reprogramming system, senescent cells secret IL-6 to promote nearby cell reprogramming (Mosteiro et al., 2016; Mosteiro et al., 2018), but the senescent cells themself are inefficient in reprograming (Banito et al., 2009; Hanna et al., 2009; Hong et al., 2009; Kawamura et al., 2009; Li et al., 2009; Marion et al., 2009; Park et al., 2008; Utikal et al., 2009; Zhao et al., 2008). Whether YAP plays a role in these processes and whether CYR61’s mechanism of action involves inflammation or senescence requires further investigation.

Outside of pluripotency, YAP is well-known for its role in controlling organ size (Camargo et al., 2007; Lee et al., 2016; Richardson and Portela, 2017; Yimlamai et al., 2014). YAP deregulation not only results in overgrown tissue and organs (i.e., the “Hippo”; phenotype), but also has a well-documented role in cancer (Harvey et al., 2013). While this has been traditionally attributed to YAP’s cell-autonomous role, our work suggests the possibility that deregulated YAP promotes tissue overgrowth and malignancy in part by altering the local secretory microenvironment or the tumor niche, a notion that is supported by a concurrent manuscript by Mugahid et al. Cancer associated fibroblasts (CAFs) and tumor associated macrophages (TAMs) are reasonable candidate cell types in this context, as their involvement in cancer by modulating the secretory microenvironment is well-documented (Aras and Zaidi, 2017; Kalluri, 2016). YAP’s non-cell-autonomous role in tumor development should be examined next.

## MATERIALS AND METHODS

### Mice, plasmid constructs

All mouse work was approved by the Institutional Animal Care and Use Committee (IACUC) of Yale University. All research animals were housed and maintained in facilities of Yale Animal Resource Center (YARC). Plasmids were generated using the Invitrogen Gateway system. The reprogramming factors OSKM and mCherry reporter are fused with 2A, and cloned into the pFUW lentiviral backbones. YAP-GFP construct was generated by PCR amplifying YAP-GFP from pEGFP C3-YAP-deltaC (Marius Sudol, Addgene # 21126) with Gateway-compatible primers to then insert into a Gateway-compatible lentiviral backbone containing a doxycycline-inducible (tetO) promoter FU-tetO-Gateway-PGK-puro. Similarly, wtYAP-mCherry and caYAP-mCherry constructs were generated by PCR amplifying the YAP sequence from YAP-GFP or caYAP from pQCXIH-Glag-YAP-S127/381A (Kunliang Guan, Addgene # 33069), respectively, and fusing to PCR-amplified mCherry sequence from OSKM-mCherry. Plasmid encoding Cre recombinase was a gift from the Valentina Greco lab. *Oct4:GFP* × *Rosa26:rtTA* mice were derived as previously described (Guo et al., 2014). Reprogrammable mice *(Col1a1:OKSM; Oct4:GFP)* (Stadtfeld et al., 2010) were purchased from the Jackson Laboratory and crossed with *Rosa26:rtTA* mice to achieve doxycycline-inducible factor expression. YAPKI and YAPKO mice were a gift from Ruslan Medzhitov and previously described and characterized (Su et al., 2015), and crossed with *Rosa26:rtTA* and (*teto)*_*7*_*-Cre* mice (Perl et al., 2002) to achieve doxycycline-inducible Cre-mediated recombination for the induction or deactivation of YAP expression, respectively. Embryonic stem cells from YAPKI or Oct4:GFP strains were derived from embryonic day 3.5 blastocysts at Yale Animal Genomic Services. Tail DNA was used for genotyping by PCR using Direct PCR Lysis Reagent (Viagen). Upon thawing, cells were plated onto gelatinized with 0.1% gelatin (American Bio) for 15 minutes at room temperature to support adhesion.

### MEF culture

Primary MEFs were isolated from day 13.5 embryos as previously described (Takahashi and Yamanaka, 2006) and cultured in filter-sterilized (0.22 μm vacuum filter; Corning) MEF medium: Dulbecco’s Modified Eagle Medium containing 4.5g/L D-Glucose, L-Glutamine and 110mg/L Sodium Pyruvate (DMEM 1X; Gibco), supplemented with heat-inactivated (30 minutes at 56^°^C) fetal bovine serum (Performance Plus FBS, United States origin; Gibco), and penicillin streptomycin glutamine at a final concentration of 100units/mL penicillin/streptomycin and 0.292mg/mL L-Glutamine (PSG 100X; Gibco). During reprogramming, cells were cultured in filter-sterilized (0.22 μm vacuum filter; Corning) ESC medium: DMEM with 15% ESC-Qualified FBS (EmbryoMax FBS; Millipore) supplemented with PSG (Gibco) in addition to non-essential amino acids (NEAA; Gibco), *β*-mercaptoethanol (BME; American Bio), and leukemia inhibitory factor (LIF; 1000U/mL EMD Millipore). Doxycycline (Sigma) was used at a concentration of 2 μg/mL when indicated. MEFs were cultured at 37^°^C/5% CO_2_.

### GMP harvest and culture

Primary GMPs are isolated from mice bone marrow located in the tibiae, femur, and iliac crest of the leg bones. All muscle and tissue was removed and the bones are kept in Dulbecco’s PBS supplemented with 2% heat-inactivated (30 minutes at 56^°^C) FBS (Gibco). Harvested leg bones were crushed with a mortar and pestle, and the resulting liquid is filtered through a 70 μm cell strainer (BD Falcon). The bone marrow was stained with a mixture of biotinylated anti-mouse antibodies to Mac-1α, GR-1, CD3, CD4, CD8a, B220, and Ter119 (BD Biosciences) for 15 minutes at 4^°^C for the depletion of the bone marrow of unwanted lineages. Cells were then washed in 2% FBS PBS, spun down into pellet (1500 rpm for 5 min), and re-suspended into streptavidin-conjugated magnetic beads (BD Biosciences) for 15 minutes at 4^°^C. Once again, cells were washed in 2% FBS PBS, spun down, and re-suspended in 3 mL 2% FBS PBS before applying onto a magnetic column (BD Biosciences) through a 70 μm basket cell strainer (BD Falcon). Flow through was then centrifuged at 1500 rpm for 5 minutes and re-suspended in an antibody cocktail (BD Biosciences) in 2% FBS PBS containing conjugated-antibodies against Kit-APC (1:100), Sca-PE (1:300), SA-BV150 (1:300), CD34-AF700 (1:20), and CD16/32-PE-Cy7 (1:150) for 15 minutes at room temperature. Stained cells were then washed and spun down for a final time at 1500 rpm for 5 minutes before re-suspending in PBS and filtering through a FACS tube with a cell strainer cap (BD Falcon). GMPs were then sorted on a BD FACS Aria through a 80 μm nozzle into 2% FBS DMEM collection medium using the marker designation Lin^−^Kit^+^Sca^−^ CD34^+^CD16/32^+^. After sorting, reprogrammable GMPs were directly cultured in ESC medium containing doxycycline on irradiated feeder MEFs, while GMPs that required viral introduction of the reprogramming factors were lentivirally-transduced overnight in a 96-well plate coated with retronectin (Takara) in the presence of concentrated virus, 5 μg/mL polybrene (EMD Millipore), and hematopoietic growth factors (PeproTech: 100ng/mL mSCF, 50 ng/mL mIL3, 50 ng/mL Flt3L, and 50 ng/mL mTPO). GMPs were cultured at 37^°^C/5% CO_2_.

### iPSC/ESC culture

Mouse embryonic stem cells were cultured on a layer of inactivated feeder MEFs at a cell density of 4.5 × 10^4^ feeder MEFs/cm^2^. Pluripotent cells were cultured in ESC medium as described above for reprogramming MEF culture. Proper maintenance of PSCs requires frequent passages upon semi-confluency, approximately every other day, onto a fresh layer of feeder MEFs, plated on the previous day at about 4.5 × 10^4^ feeder MEFs/cm^2^. Pluripotent stem cells were cultured at 37^°^C/5% CO_2_.

### Generation of wildtype, EV-, and YAP-feeder MEFs

Primary MEF cells were harvested from day 13.5 embryos as previously described (Takahashi and Yamanaka, 2006) and expanded over maximally four passages. During each passage, cells are trypsanized, pooled, spun down, and replated in MEF medium so as to achieve a 1:3 or 1:2 expansion. Cells were then irradiated using gamma irradiation at 8000 rads. After irradiation, cells were frozen down in manageable aliquots and stored in liquid nitrogen. Cells were thawed as needed and plated at a cell density of 4.5 × 10^4^ feeder MEFs/cm^2^. For EV- and YAP-feeder MEFs, primary MEFs were first transduced with lentiviral constructs encoding EV or YAP, and then sorted based upon mCherry fluorescence and replated for expansion before irradiation.

### Lentivirus production and transduction

Lentivirus was prepared from 293T cells in 10cm dishes cultured in MEF medium. 293Ts were plated at 7 × 10^6^ cells/10cm plate the night before transfection. FuGENE6 (Promega) was used to mediate efficient entry of plasmid mixture containing 11 μg of the appropriate lentiviral construct, 5.5 μg of VSV-G/pMD2.G, and 8.25 μg of pCMV-DR8.91 into 293Ts. The day following transfection, the medium was changed to high BSA medium: DMEM with 10% heat-inactivated (30 minutes at 56^°^C) fetal bovine serum (Performance Plus FBS, United States origin; Gibco) plus 1% PSG (Gibco) and 1.1g/100mL BSA (American Bio), sterilized using 0.22 μm Corning vacuum filters). Each day, the medium was collected and fresh high BSA medium was added. After three days of collection, the viral collections were pooled and spun down for 5 minutes at 1500 rpm before either using directly as fresh virus or concentrating in a viral pellet at 20,000 rpm for 90 minutes at 4^°^C. For transduction, virus is added to cells at final concentration of 5 μg/mL polybrene (EMD Millipore) overnight.

### Co-culture experiments

Using the ibidi co-culture device, cells were plated within the individual minor wells, as discussed. Primary GMPs were isolated from the reprogrammable mouse model (Stadtfeld et al., 2010). A fixed number of GMPs were sorted into individual wells of a 96-well plate via a BD FACS Aria machine, to be re-suspended into ESC medium containing doxycycline to achieve a final plating of twenty-five GMPs per central minor well on top of a layer of wildtype feeder MEFs. EV-feeders and YAP-feeders were plated in the outer wells surrounding the central minor well (excluding the two wells immediately to the left and right of the minor well) at a density of 4.5 × 10^4^ feeder MEFs/cm^2^. The following morning, ESC medium containing doxycycline was carefully added to the entire major well. Cells were cultured without medium change for five days of reprogramming.

### Conditioned medium collection and cytokine/growth factor identification assay

For the preparation of conditioned medium, EV- and YAP-feeders were first plated at a density of 4.5 × 10^4^ feeder MEFs/cm^2^ in ESC medium containing doxycycline. Once feeders become adherent, reprogrammable GMPs were then harvested and plated on top of the feeder layer. After 24 hrs, the medium was collected and centrifuged to remove cells and cellular debris, before adding directly to reprogramming cultures. When indicated, conditioned medium was diluted with fresh medium containing doxycycline to achieve 10%, 25%, 50%, and 75% conditioned medium. For cytokine/growth factor identification, medium conditioned for 5 days within the co-culture device (Ibidi) was collected and centrifuged before assaying directly with the Proteome Profiler Mouse XL Cytokine Array Kit (R&D Systems). Spot densitometry was quantified using the QuickSpots software (H&L Image).

### Reprogramming of MEFs and GMPs from reprogrammable mouse system

Primary MEFs and GMPs were isolated from the reprogrammable mouse model conditionally expressing Oct4, Klf4, Sox2, and c-Myc (OKSM) under a doxycycline-inducible promoter, in the context of the m2 reverse tetracycline-controlled transactivator (rtTA) (Stadtfeld et al., 2010). Upon the first passage after harvest, MEFs were seeded at a cell density of 1 × 10^4^ cells/cm^2^ for overnight transduction. Primary GMPs were harvested as previously described (above) and plated in a 96 well for overnight transduction. MEFs and GMPs were transduced overnight with 5 μg/mL polybrene transfection reagent (EMD Millipore) and either unconcentrated viral supernatant or the appropriate culture medium containing concentrated virus. As indicated, MEFs were sorted using BD FACS Aria based on fluorescent protein reporters encoded by the viral constructs. After sort, MEFs were re-seeded at a cell density of 1 × 10^4^ cells/cm^2^ on top of a previously plated MEF feeder layer of 4.5 × 10^4^ cells/cm^2^. MEFs were then allowed to reprogram, with regular medium changes (every 2-3 days) with fresh ESC medium and 2 μg/mL doxycycline to sustain the expression of transgenic OSKM. Reprogrammable GMPs were replated into fresh ESC medium plus 2 μg/mL doxycycline after sort and/or transduction at a concentration of 1 × 10^3^ cells/3.5cm^2^ on a feeder of inactivated feeder MEFs.

### Calculation of Reprogramming Efficiency

Equal numbers of reprogramming cell cultures were plated in the presence of doxycycline on top of irradiated feeder MEFs. Efficiencies were determined after colony formation by dividing the number of Oct4:GFP+ or Nanog+ iPSC colonies by the starting number of seeded cells.

### RNA preparation and analysis

Total RNA was extracted with Trizol^®^ reagent and reverse transcribed (Invitrogen). After homogenization, the RNA was purified and prepared according to the Invitrogen Trizol protocol. Using the Invitrogen SuperScript III First-Strand Synthesis System for RT-PCR, RNA was reverse transcribed into cDNA for downstream analysis by quantitative real-time PCR. Quantitative real-time PCR was performed using the iQ™ SYBR^®^ Green Supermix (Bio-Rad) on a Bio-Rad CFX96. For high-throughput RNA-sequencing performed with the Illumina HiSeq 2000 Sequencing System, total RNA was used directly for RNA library preparation, following the manufacturer’s instructions. Prior to initiating sequencing procedure, the quality of total RNA was analyzed on Agilent Bioanalyzer. The RNA sample that has more than 8 RNA integration number (RIN) was taken up for seq library preparation using TruSeq Stranded mRNA Library Preparation Kit supplied by Illumina (Cat # RS-122-2101). The protocol followed was as per manufacturer’s instruction. For data analysis, RNA-seq reads were aligned to the mouse genome mm10 using Tophat followed by differential gene expression analysis with Cufflinks in Galaxy.

### Immunofluorescence and image acquisition

For immunofluorescence, cells were washed three times with PBS, fixed with 4% paraformaldehyde (diluted from 37% solution from Sigma) for 15 minutes at room temperature, washed three times with PBS and permeabilized with 0.25% Triton X-100 (American Bio) solution in PBS for 10 minutes at room temperature. Samples were then blocked for 1 hr at room temperature with rocking with AbDil buffer: 0.1% sodium azide (American Bio), 0.1% Triton X-100 (American Bio), 2% BSA (American Bio) in 1X PBS, stored at 4^°^C). Cells were then stained overnight at 4^°^C with rocking in primary antibodies diluted in AbDil buffer against mouse NANOG and YAP (Cell Signaling, 1:500) and CYR61 (Cell Signaling, 1:500). The cells were then washed three times for five minutes each in AbDil and incubated with the anti-rabbit secondary antibody conjugated with Alexa Fluor 680 (Cell Signaling, 1:500) for 1 hr at room temperature and rocking. DNA was stained with DAPI (Invitrogen, 1:1000). Images were taken with the Leica SP5 confocal microscope, Olympus IX51 inverted fluorescence microscope, a LAXCO LMI-6000 fluorescence microscope, or the ImageExpress Micro 4 high-content imaging system (Molecular Devices) paired with a PC image analysis station equipped with MetaXpress software (Molecular Devices).

### FACS analysis and sorting

For flow cytometry, cells were directly flowed after trypsanization (i.e., no antibody staining), as they contained intrinsic fluorescent reporters, e.g., mCherry or GFP. Cells were washed in PBS twice before being trypsanized in 0.25% Trypsin-EDTA (Gibco) and collected in serum-containing medium to quench trypsin activity before centrifugation at 1500 rpm for 5 minutes. Cell pellets were then washed and re-suspended in PBS and filtered through a 40 mm cell strainer cap FACS tube (BD Falcon) to achieve single-cell suspension. With the exception of GMP isolation from bone marrow by conjugated antibodies (as described above), FACS was carried out using intrinsic or virally-encoded fluorescent reporters and, thus, could be carried out directly after trypsanization without antibody-staining. Cells were sorted on BD FACS Aria and/or analyzed on a BD LSRII. Flow data was analyzed using FlowJo software.

### Immunoblot

Cell lysates were harvested by directly lysing the 2 × 10^^6^ cells with 2x sampling buffer (Bio-Rad). Proteins were separated by SDS-PAGE, transferred onto nitrocellulose membranes (Bio-Rad). The membranes were blocked with 10% nonfat dry milk in TBS-Tween (TBST) for 1 hour, incubated with primary antibodies overnight at 4^°^C, followed by incubation with horseradish-peroxidase-conjugated secondary antibodies for 1 hour, and illuminated by enhanced chemiluminescence (ECL).

### AP staining

AP staining was performed using the AP staining kit from StemGent (00-0055).

**Supplemental Table 1:**
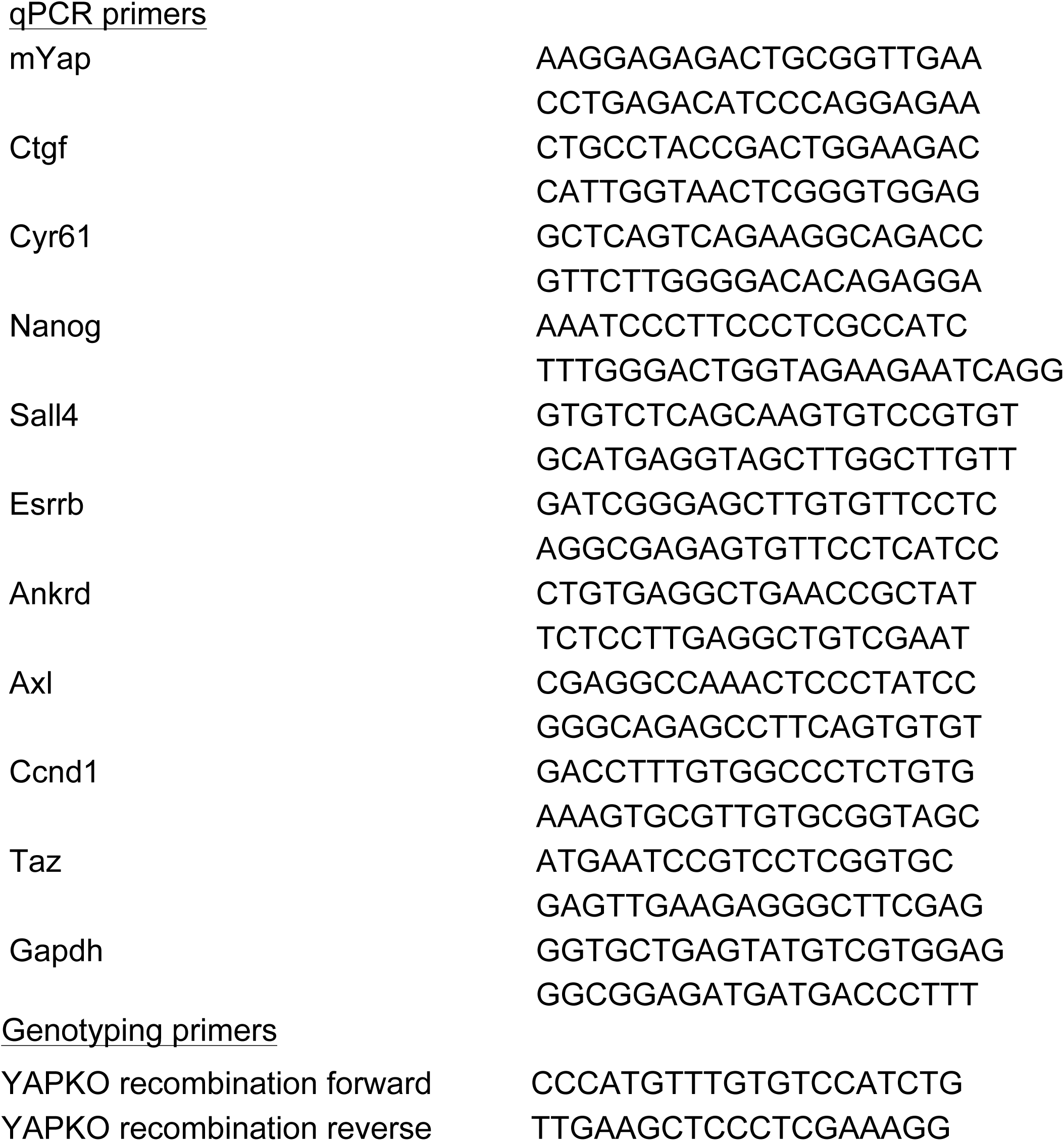
Primer sequences

---

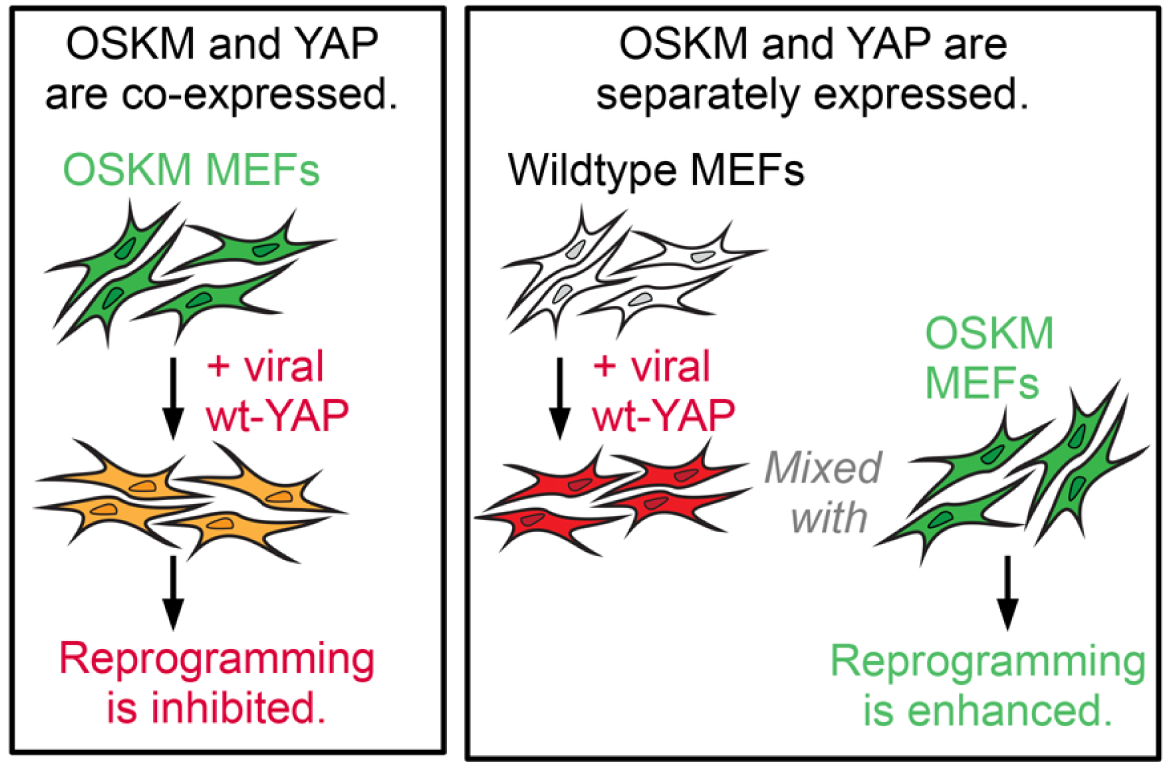

**Figure S1:**
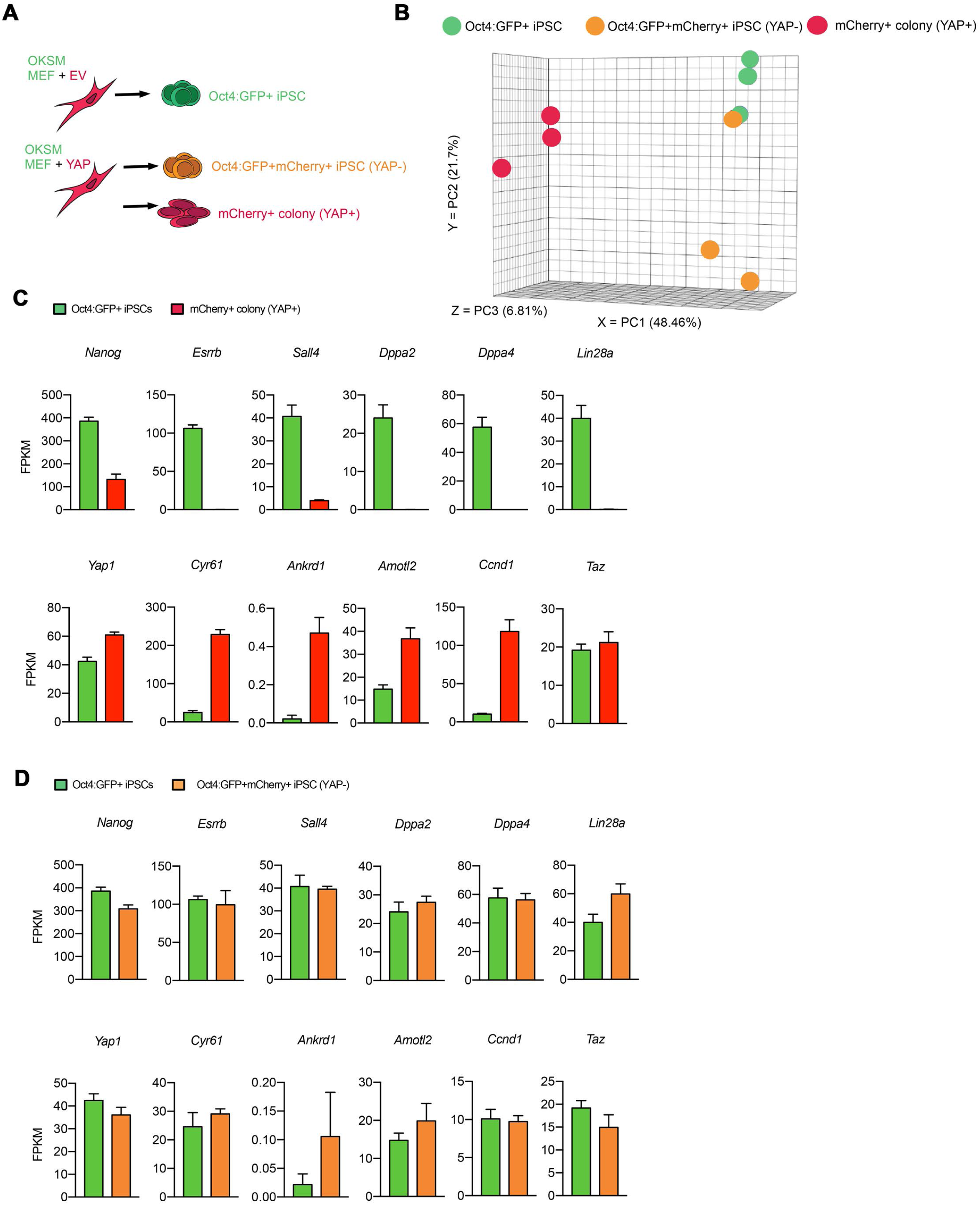
YAP inhibits pluripotency induction cell-autonomously from MEFs. (A) Experimental scheme illustrating the three cell types analyzed, following the same procedures shown in Fig 1A. These include the Oct4:GFP and mCherry double-positive cells, mCherry+ flat colony-forming cells from YAP-transduced cultures, and Oct4:GFP+ iPSCs from EV control cultures. (B) Principle Component Analysis (PCA) plot of the three cell types illustrated in (A) analyzed by mRNA-seq. (C) Expression levels of endogenous pluripotency network genes *Nanog*, *Esrrb*, *Sa/14*, in addition to *Yap* pathway genes *Cyr61*, *Akrd1*, *Amot/2*, *Ccnd1*, and *T az* by mRNA-seq. Comparison is made between control Oct4GFP+ iPSCs and the mCherry+ flat colony forming cells from YAP-transduced cultures. (D) Similar to (C), comparison is made between control Oct4GFP+ iPSCs and the Oct4:GFP+ cells that remained mCherry+ from YAP-transduced cultures.

**Figure S2:**
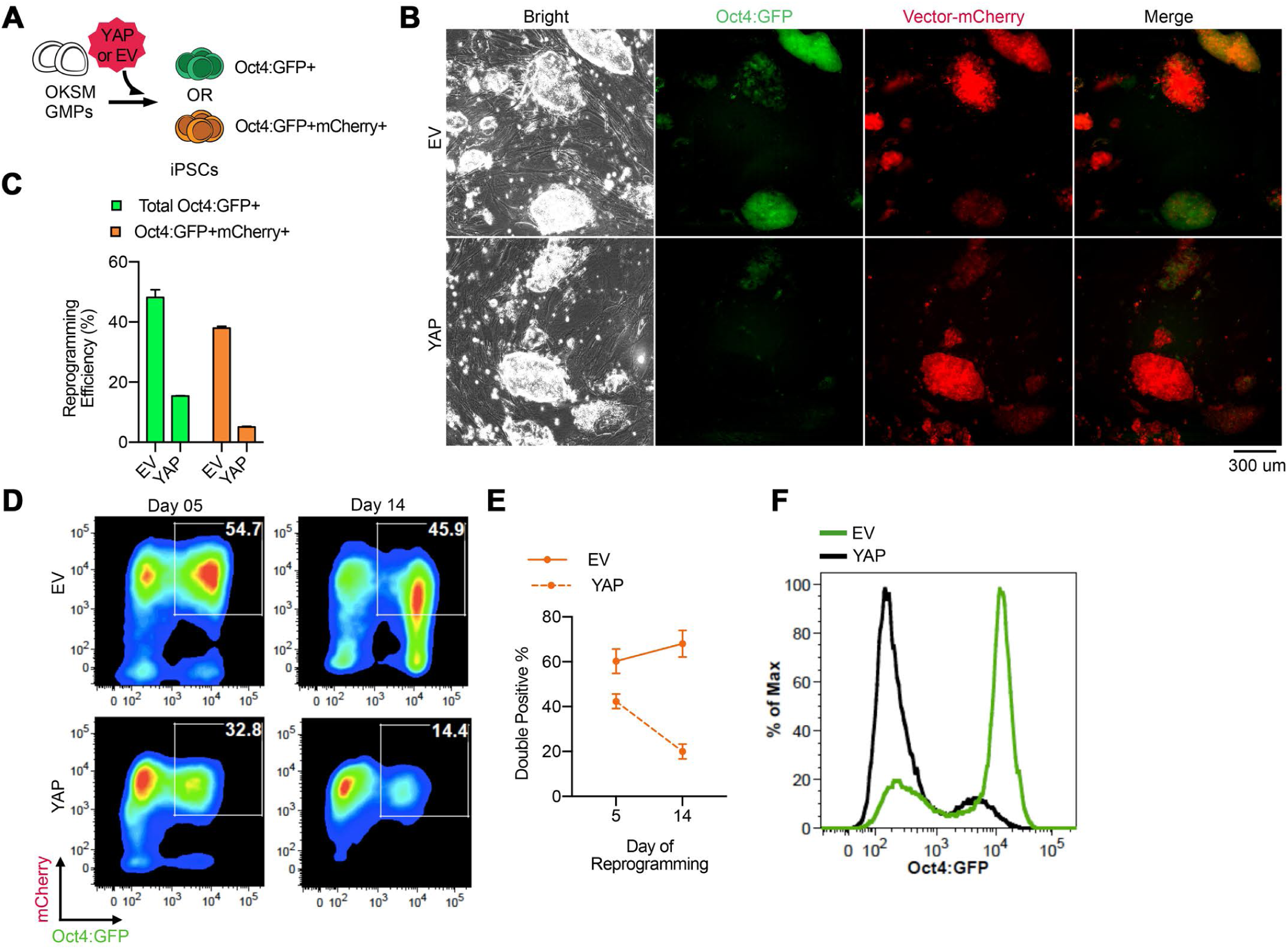
YAP inhibits pluripotency induction cell-autonomously from hematopoietic progenitors. (A) Experimental scheme illustrating OKSM GMPs transduced with viral vectors as shown in Fig 1A. The expression status of Oct4:GFP was determined in the resulting iPSCs in relation to their expression of mCherry, shown in (B-F). (B) Representative images of the resulting colonies derived from reprogrammable GMPs transduced with EV or YAP. (C) Quantification of reprogramming efficiency based on total Oct4:GFP+ colonies or Oct4:GFP and mCherry double-positive colonies. (D) Representative FACS plots of reprogrammable GMPs transduced with EV or YAP on day 5 and day 14 of reprogramming. (E) Quantification of the percentage of Oct4:GFP and mCherry double-positive cells, on day 5 and day 14 of reprogramming. (F) Oct4:GFP fluorescence intensity on day 14 of reprogramming.

**Figure S3:**
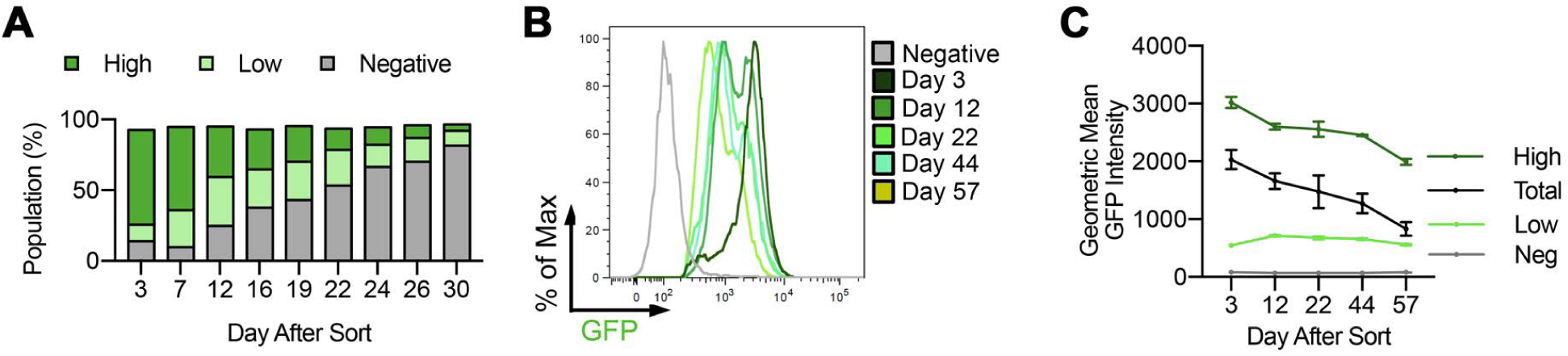
Ectopic YAP expression does not promote pluripotency maintenance. (A) Quantification of the GFP+ populations in YAPKI mESC cultures following Cre-mediated recombination and FACS-sort. Days indicate time after FACS-sort. (B) Representative FACS plot showing the evolution of fluorescence intensity of the GFP+ population over time. (C) Quantification of GFP geometric mean fluorescence intensity in the total GFP+ population (Total), GFP-low (Low), GFP-high (High) and GFP-negative (Negative) populations over time.

**Supplemental Figure 4:**
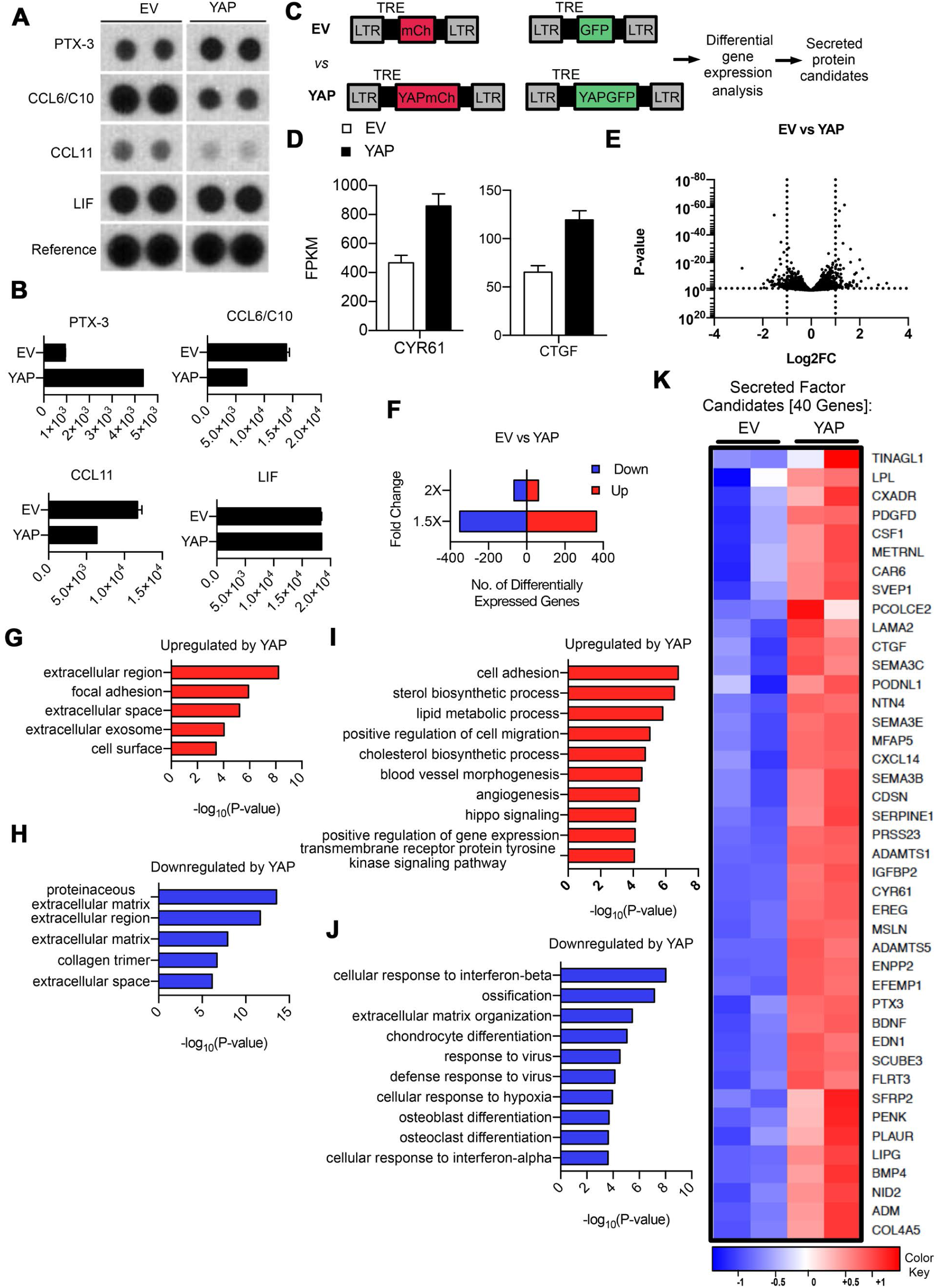
YAP alters the expression of microenvironmental proteins. (A) Cytokine array spot densities of detected proteins in the conditioned medium from EV and YAP feeders, harvested from co-culture devices as shown in Fig 5D. (B) Quantification of the spot density shown in (A). (C) Schema of the viral constructs used to generate MEFs expressing the control vectors (EV for GFP or mCherry) or YAP fused to GFP or mCherry, which were analyzed by mRNA-seq. The differentially expressed genes were further analyzed in (D-K). (D) mRNA-seq data validating elevated YAP target genes *Cyr61* and *Ctgf* in YAP-expressing MEFs. (E) Volcano plot comparing the gene expression of MEFs expressing EV or YAP. Dotted lines demarcate genes with significant differential expression of more than 2-fold and with a p-value of < 0.05. (F) Number of genes that are upregulated or downregulated by 1.5-fold or 2-fold between EV and YAP-expressing MEFs. (G) Upregulated cellular components in YAP-expressing MEFs by Gene Ontology (GO) analysis. (H) Downregulated cellular components in YAP-expressing MEFs by GO analysis. (I) Upregulated pathways in YAP-expressing MEFs by GO analysis. (J) Downregulated pathways in YAP-expressing MEFs by GO analysis. (K) Heatmap of genes encoding candidate secreted proteins that are upregulated at least 2-fold in YAP-expressing MEFs.

**Figure S5:**
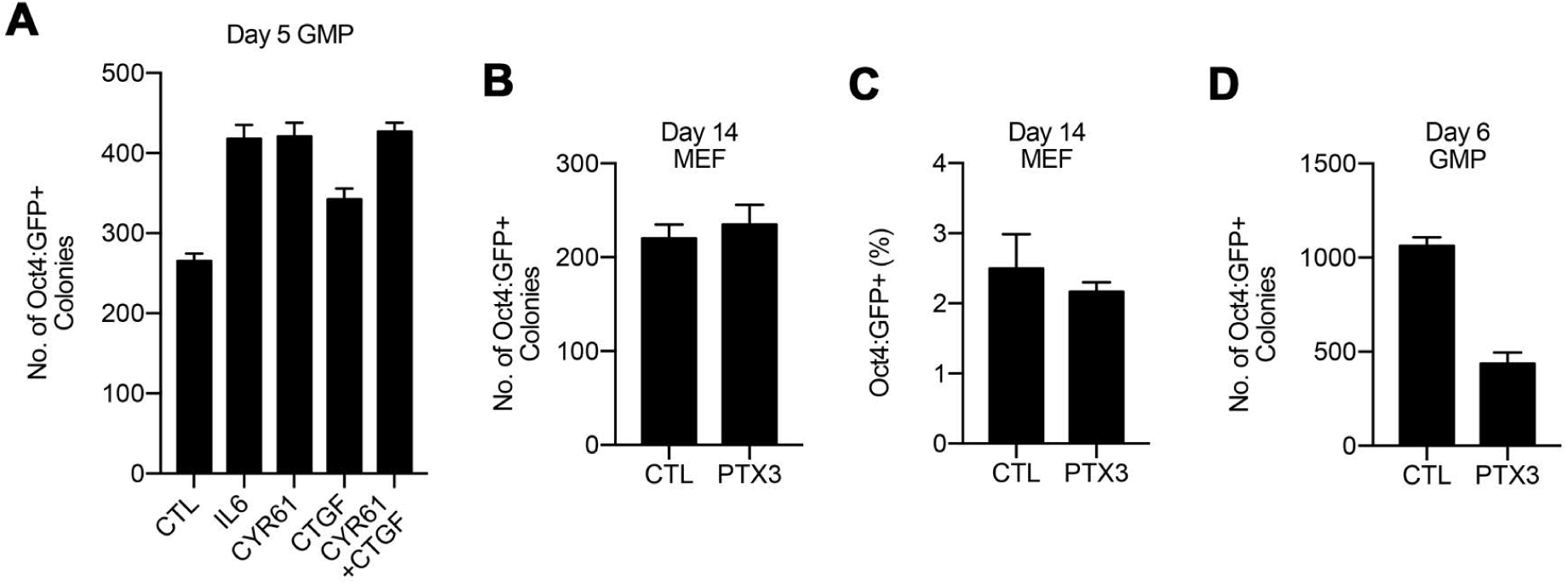
Recombinant CYR61 protein promotes pluripotency induction. (A) Quantification of the number of Oct4:GFP+ colonies from reprogrammable GMPs on day 5 of reprogramming, in the presence of various recombinant proteins. (B) Quantification of the number of Oct4:GFP+ colonies from reprogrammable MEFs on day 15 of reprogramming in the presence of recombinant Pentraxin-3. (C) Quantification of the percent of Oct4:GFP+ from reprogrammable MEFs on day 15 of reprogramming in the presence of recombinant Pentraxin-3 Quantification of the number of Oct4:GFP+ colonies from reprogrammable GMPs on day6 of reprogramming in the presence of recombinant Pentraxin-3.

**Figure S6:**
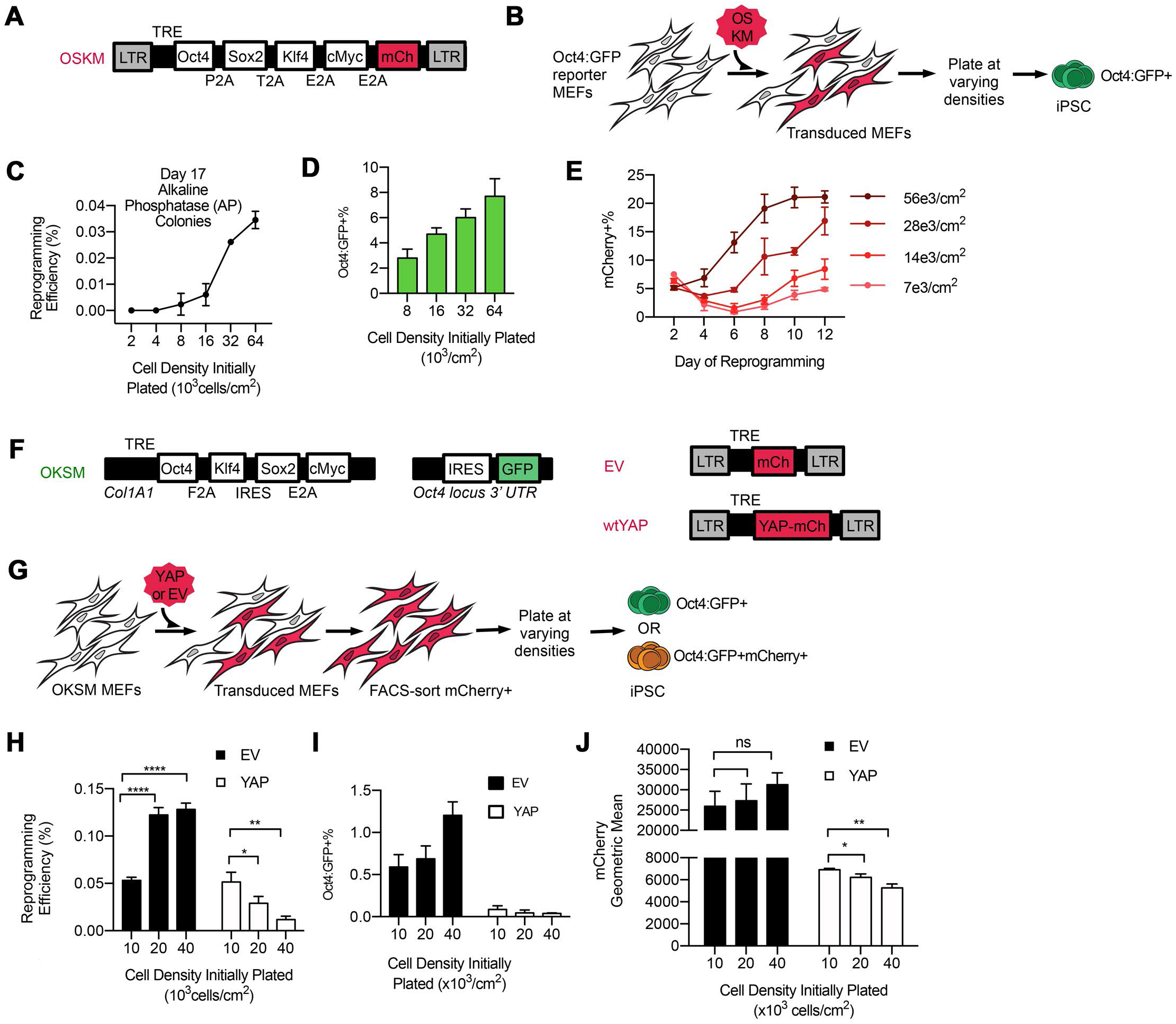
Cell plating density modulates reprogramming efficiency. (A) Schema of polycistronic lentiviral vector encoding OSKM-mCherry, similar to Fig 2A and 4A. (B) Experimental scheme illustrating MEFs, which express the Oct4:GFP reporter, were transduced with OSKM-mCherry, trypsinized and replated at varying cell densities to allow further reprogramming. The results are shown in (C-E). (C) Quantification of reprogramming efficiency for cells plated at different densities based on alkaline phosphatase (AP) positive colonies on day 17 of reprogramming. (D) Quantification of the percent of Oct4:GFP+ cells which emerge from cultures plated at different densities on day 18 of reprogramming. (E) Quantification of the percent of mCherry+ cells in cultures plated at different densities throughout reprogramming. (F) Schema of lentiviral constructs for transducing EV or YAP into reprogrammable MEFs expressing transgenic OKSM and Oct4:GFP reporter, similar to Fig 1A-B. (G) Experimental scheme illustrating reprogrammable MEFs co-expressing EV or YAP which were FAGS-sorted and plated at varying densities to allow further reprogramming. The results are shown in (H-J). (H) Quantification of reprogramming efficiency based on Oct4:GFP+ colonies on day 15 of reprogramming. (I) Quantification of the percent of Oct4:GFP+ cells in cultures plated at varying densities on day 12 of reprogramming. (J) The geometric mean of mCherry fluorescence in cultures plated at varying densities expressing EV or YAP on day 12 of reprogramming.

